# A new haplotype-resolved turkey genome to enable turkey genetics and genomics research

**DOI:** 10.1101/2022.08.18.504375

**Authors:** Carolina P. Barros, Martijn F.L. Derks, Jeff Mohr, Benjamin Wood, Richard P.M.A. Crooijmans, Hendrik-Jan Megens, Marco C.A.M. Bink, Martien A.M. Groenen

**Affiliations:** Wageningen University and Research, Wageningen, Netherlands; Hybrid Turkeys, Kitchener, ON, Canada; School of Veterinary Science, University of Queensland, Gatton, QLD, Australia; Hendrix Genetics Research, Technology & Services, Boxmeer, Netherlands

**Keywords:** Genome assembly, Turkey genomics, trio-binning, animal breeding

## Abstract

**Background:** The domesticated turkey (*Meleagris gallopavo*) is a species of significant agricultural importance and is the second largest contributor, behind broiler chickens, to world poultry meat production. The previous genome is of draft quality and partly based on the chicken (*Gallus gallus*) genome. A high-quality reference genome of *Meleagris gallopavo* is essential for turkey genomics and genetics research and the breeding industry.

**Results:** By adopting the trio-binning approach, we were able to assemble a high-quality chromosome-level F1 assembly and two parental haplotype assemblies, leveraging long-read technologies and genomewide chromatin interaction data (Hi-C). These assemblies cover 35 chromosomes in a single scaffold and show improved genome completeness and continuity. The three assemblies are of higher quality than the previous draft quality assembly and comparable to the current chicken assemblies (GRCg6a and GRCg7). Comparative analyses reveal a large inversion of around 19 Mbp on the Z chromosome not found in other Galliformes. Structural variation between the parent haplotypes were identified in genes involved in growth providing new target genes for breeding.

**Conclusions:** Collectively, we present a new high quality chromosome level turkey genome, which will significantly contribute to turkey and avian genomics research and benefit the turkey breeding industry.

## Introduction

The domesticated turkey (*Meleagris gallopavo*) is an important agricultural species and the second largest contributor to world poultry production [1]. The turkey is a member of the Phasianidae family within the order Galliformes. Turkeys and chickens diverged about 25-40 million years ago [2]. Despite the relative long divergence time, the genome synteny and karyotype of both are highly conserved (Griffin et al., 2007). The turkey has 2n=80 compared to the chicken with 2n=78. Most of the chromosomes are small microchromosomes, while only a few macrochromosomes are present in the karyotype. The turkey karyotype is very similar to the chicken, except that chicken chromosome 2 is homologous to two turkey chromosomes (chromosomes 3 and 6) and chicken chromosome 4 is homologous to turkey chromosomes 4 and 9 [4]. In addition, a high degree of synteny has also been observed between the chicken and turkey genomes [5].

The first turkey genome assembly (UMD2), published in 2010, was among the first to be done almost exclusively based on second generation sequencing data, and by current standards would be considered of draft quality [5]. The authors produced a chromosome level assembly and assembled 30 autosomal and two sex chromosomes. The assembly included linkage data based on a low-density genetic map and the placement of scaffolds to chromosomes relied considerably on conserved synteny assumptions with the better assembled chicken (*Gallus gallus*) genome. However, that version of the chicken genome had many microchromosomes missing altogether or only partially characterized. Avian microchromosomes have proved to be difficult to assemble even today. Reliance on an incomplete chicken genome and the general difficulty in assembling the avian microchromosomes resulted in a poor representation of microchromosomes in that first UMD2 turkey genome. An updated version of the turkey genome (Turkey_5.1; GCA_000146605.4) has been available since 2019, though it still shows low gene completeness and an incomplete set of microchromosomes.

The problems in characterizing microchromosomes are partly due to sequence characteristics, i.e., high GC and repeat content in microchromosomes, and partly due to their extremely small size and lack of genetic linkage group markers to differentiate the microchromosomes from other chromosomes [6]. Ongoing efforts in producing high quality assemblies of the microchromosomes in avian genomes have been unsuccessful due to multiple causes [6].

High quality genome sequences are an essential resource for research and applications in the life sciences. In domestic animal breeding, genome wide marker panels are routinely used to support genomic selection and this significantly accelerates genetic progress[7]. An improved genome sequence facilitates ongoing genomic breeding programs. Furthermore, an improved genome assembly will greatly enhance functional interpretation of genomic variation in those breeding populations. For instance, improved annotation of (non)-coding genes benefits the functional interpretation of genome wide association studies (GWAS), and aids in identifying targets for gene editing [8].

Currently, more species in the Galliformes have high quality long-read based assemblies, including the Japanese quail [9]. Gunnison sage-grouse [10], and the helmeted guineafowl [11], allowing for comparative studies within the Galliformes. The genome assemblies of turkey (this paper) and chicken, however, are of considerably higher quality compared to other Galliforme species. This provides opportunities for an in-depth comparison between the two most important avian agricultural species.

Third generation sequencing techniques have made it possible to produce high quality chromosome-based assemblies. The chicken GRCg6a assembly and more recently individual broiler (GRCg7b) and layer (GRCg7w) assemblies have been produced from long read sequencing techniques. The GRCg7 genomes now include (parts of) all microchromosomes. These new chicken assemblies show superior metrics of quality and completeness to previous genome assemblies. In this study we use a relatively new technique, the trio-binning approach, to construct high quality haplotype-resolved turkey assemblies [12]. A similar approach was also applied to create the GRCg7 chicken genome assemblies. Short reads from each parent are used to resolve the F1 long reads into groups of long reads belonging to each parent. Each haplotype is then assembled independently resulting in three high quality genome assemblies, one from both parental haplotypes, and one F1 assembly. This approach is especially powerful to assess structural variation between the parental haplotypes and works well with high heterozygosity rates as this aides in the resolution of the parent haplotypes in the F1 assembly.

In this study our aims were to use the trio-binning approach to produce a chromosome-level turkey assembly (F1), and two parental haplotype assemblies. We further aim to compare the two parental haplotypes to identify structural differences. A good reference genome is essential for many research and commercial applications. In this study we highlight how our new turkey genome can benefit both research and the breeding industry.

## Results

### Data and assembly of Mgal_WUR_HG_1.0

Three individual turkeys (two parents and one F1) were sequenced using the trio binning approach [12]. The two parental animals derive from two distinct commercial lines from the breeding company Hybrid Turkeys, a Hendrix Genetics company. The F1 animal was sequenced with a depth of 270x using PacBio single-molecule real-time (SMRT) sequencing technology. Approximately 12.25 million subreads were produced with a mean length of 22.5 kb, and N50 read length of 32.5 kb. Reads were assembled using wtdgb2 assembler [13] resulting in an initial assembly comprising of 315 contigs with an N50 of 26.68 Mb. The assembly was further scaffolded using Hi-C with HiRise [14]. Additional scaffolding was performed using SALSA (with Hi-C) [15] and Redundans [16]. The scaffolded assembly was subsequently polished with short reads (three rounds) to produce a final chromosome-level assembly consisting of 151 scaffolds and 232 contigs with a scaffold N50 of 70 Mbp and contig N50 of 26.55 Mbp **(Table 1)**. This captures the chromosome arms in a single contig. The Hi-C contact map can be found in **Supplementary File** 1: **Figure S1.**

**Table 1:** Assembly statistics. Summary statistics for the new Mgal_WU_HG_1.0 and parental assemblies, and comparison with previous turkey assembly (Turkey_5.1) and recent broiler assembly (GRCg7b).

|  | Mgal_WU_HG_1.0 | Turkey_5.1 | GRCg7b | Parent 1 | Parent 2 |
| --- | --- | --- | --- | --- | --- |
| **Total sequence length (bp)** | 1,001,818,376 | 1,115,474,681 | 1,053,332,251 | 1,051,251,094 | 1,085,657,715 |
| **Length ungapped (bp)** | 1,001,806,830 | 1,080,180,254 | 1,049,948,333 | 1,050,601,018 | 1,085,166,758 |
| **No. of scaffolds** | 151 | 187,695 | 214 | 415 | 489 |
| **No. of unplaced scaffolds** | 115 | 187,662 | 172 | 379 | 453 |
| **No. of chromosomes** | 36 | 33 | 42 | 36 | 36 |
| **Scaffold N50 (bp)** | 70,339,173 | 3,898,092 | 90,861,225 | 71,046,337 | 71,481,950 |
| **Scaffold L50** | 5 | 80 | 4 | 4 | 4 |
| **No. of contigs** | 232 | 250,220 | 677 | 738 | 675 |
| **Contig N50 (bp)** | 26,554,504 | 27,076 | 18,834,961 | 9,174,806 | 19,817,032 |
| **Contig L50** | 12 | 11,318 | 18 | 34 | 13 |

#### Haplotype assemblies

As part of the trio-binning approach, both parental haplotypes were assembled with TrioCanu [12]. We were able to map 110X of the PacBio reads to parent 1 and 137X of the PacBio reads to parent 2, resulting in two parental haplotype assemblies with contig N50 of 9,174,806 bp and 19,855,975 bp for parents 1 and 2, respectively. We performed further scaffolding using LRscaf [17] and anchored the assemblies to the F1 assembly using RagTag [18]. The final statistics of the assemblies are shown in **Table 1.**

#### Assembly accuracy and completeness

The completeness and accuracy of the assemblies were assessed using BUSCO [19] and wholegenome alignments. All three assemblies contained over 96% of the expected avian and vertebrate gene sets, comparable to the GRCg6a and GRCg7b chicken genomes and covering 5.4% more gene space compared to the previous turkey genome assembly (Turkey_5.1), as shown in Table 2.

**Table 2:** Assembly completeness measured in BUSCO scores. Percentage of aligned genes for the vertebrae (n=3354) and avian (n=8338) gene set in the turkey and chicken assemblies.

|  | Mgal_WU_HG_1.0 |  | Turkey_5.1 |  | GRCg7b |  | Parent 1 |  | Parent 2 |  |
| --- | --- | --- | --- | --- | --- | --- | --- | --- | --- | --- |
|  | Avian | Vertebrate | Avian | Vertebrate | Avian | Vertebrate | Avian | Vertebrate | Avian | Vertebrate |
| Complete | 96.7 | 96.4 | 91.3 | 88.4 | 97.0 | 96.5 | 96.6 | 96.0 | 96.8 | 96.4 |
| Complete and single-copy | 96.4 | 95.9 | 91.1 | 87.9 | 96.7 | 95.7 | 94.8 | 93.9 | 94.1 | 93.2 |
| Complete and duplicated | 0.3 | 0.5 | 0.2 | 0.5 | 0.3 | 0.8 | 1.8 | 2.1 | 2.7 | 3.2 |
| Fragmented | 0.9 | 1.0 | 4.1 | 5.8 | 0.9 | 1.2 | 0.9 | 1.1 | 0.9 | 1.0 |
| Missing | 2.4 | 2.6 | 4.6 | 5.8 | 2.1 | 2.3 | 2.5 | 2.9 | 2.3 | 2.6 |

Second, sequence alignments of the F1 assembly were made to the GRCg7b chicken assembly and the Turkey_5.1 assembly (Figure 1). The alignment is highly congruent with the chicken genome **(Figure 1A),** indicating a high degree of conserved synteny. The main exception was a large ~19 Mbp inversion on the Z-chromosome (coordinates 44517565-63393669 bp). This inversion was also not present in the previous turkey build, Turkey_5.1, as seen in the alignment **(Figure 1B).** The alignment further shows that in the Turkey_5.1 assembly many contigs were placed in the wrong orientation (resulting in a “zigzag” alignment pattern).

**Figure 1:**
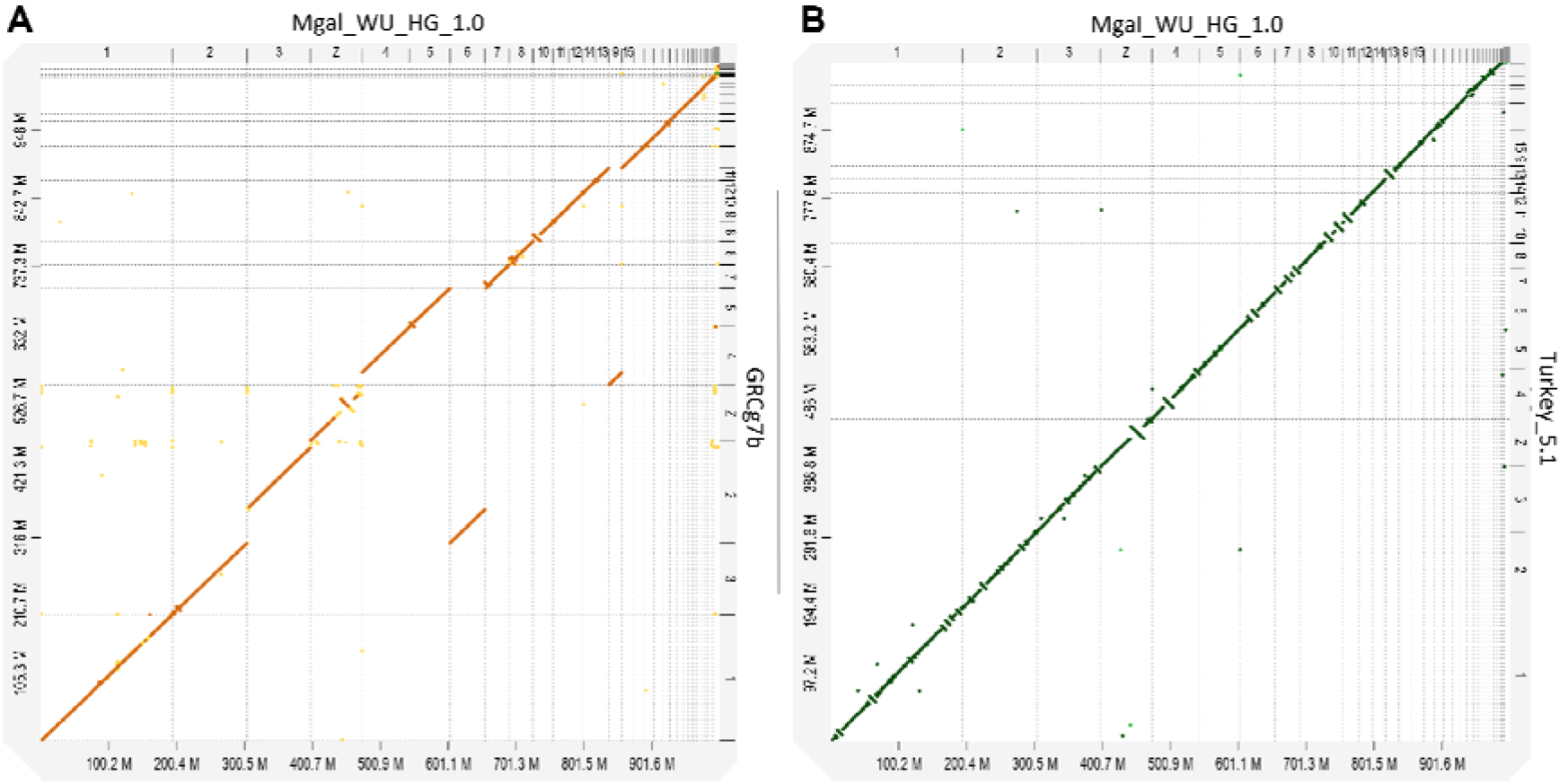
Genome-wide alignment plots. A) Mgal_WU_HG_1.0 aligned with GRCg7b. Alignment shows high structural coherence between both genomes. B) Mgal_WU_HG_1.0 aligned with the old turkey genome build Turkey_5.1. Alignment shows multiple contigs that were placed in the wrong orientation in the previous Turkey_5.1 build.

### Repeat and gene annotation

#### Repeat content

We annotated the repeats using a custom repeat library built using RepeatModeler [20]. Repeats were found to cover 10.45% of the genome. The most common were LINE elements, covering 6.35% of the genome. Furthermore, 0.76% of bases were DNA transposons, 0.53% long terminal repeats (LTRs), and 1.58% low complexity and simple repeats. The remaining 1.23% of the repeats remained unclassified.

#### Gene Annotation

The Ensembl annotation pipeline was used to annotate Mgal_WU_HG_1.0 [21]. The present annotation includes fewer annotated genes compared to Turkey_5.1 and the chicken annotations, but does include more non-coding genes, as shown in Table 3. Hence, the annotation provides a comprehensive overview of the turkey transcriptome with a large increase in transcripts compared to Turkey_5.1 and GRCg6a (Table 3). As expected, microchromosomes show higher gene density compared to macrochromosomes (Figure 2). The density generally increases with decreasing microchromosome size.

**Table 3:** Annotation statistics for the turkey (Mgal_WU_HG_1.0, Turkey_5.1) and chicken (GRCg6a, GRCg7b) genomes.

|  | Mgal_WU_HG_1.0 | Turkey_5.1 | GRCg6a | GRCg7b |
| --- | --- | --- | --- | --- |
| **Coding genes** | 16,127 | 16,226 | 16,878 | 17,007 |
| **Non-coding genes** | 7,736 | 1,585 | 7,166 | 13,040 |
| **Small non-coding genes** | 504 | 543 | 1,525 | 1,089 |
| **Long non-coding genes** | 7,228 | 1,038 | 5,506 | 11,946 |
| **Misc non-coding genes** | 4 | 4 | 135 | 5 |
| **Pseudogenes** | 45 | 159 | 312 | 61 |
| **Gene transcripts** | 53,441 | 30,708 | 39,288 | 72,689 |

**Figure 2:**
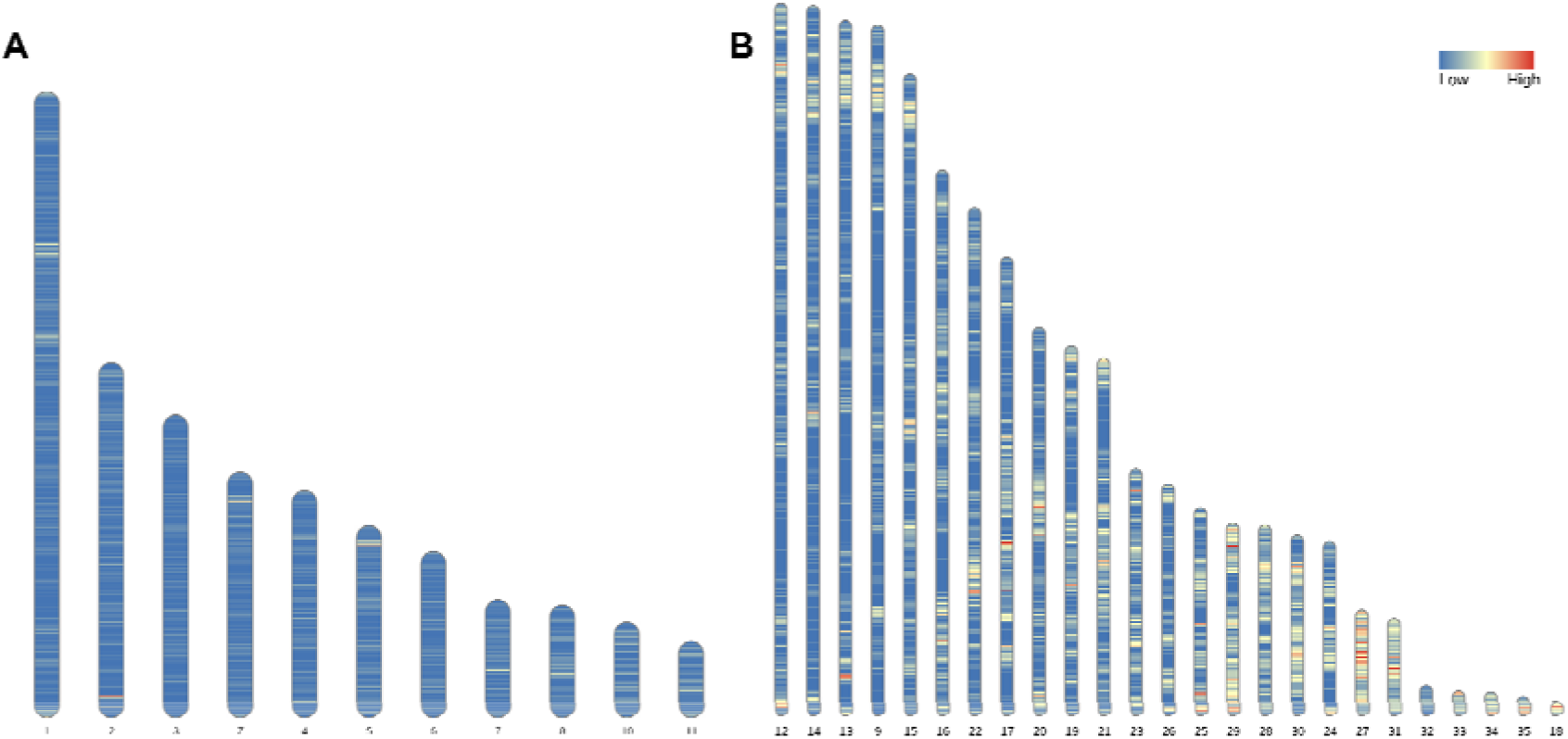
Ideogram showing gene density. A) macro (1-6, Z) and intermediate chromosomes (7,8,10,11). B) micro chromosomes (9,12-35) in the Mgal_WU_HG_1.0 genome.

We identified chicken and Turkey_5.1 homologues of the Mgal_WU_HG_1.0 genes **(Supplementary File 1: Table S1)**. The majority of the protein-coding genes have a 1:1 orthologue in the Turkey_5.1 (82.4%) or in the GRCg6a (86.3%) genome assemblies. The higher number of genes orthologous to the most recent chicken assemblies supports our assertion of a significant improvement of assembly and annotation quality compared to Turkey_5.1

#### Lineage specific expansion and contraction of protein-coding gene families

OrthoFinder [22] was used to infer orthogroups from the following set of bird species - turkey, chicken, Japanese quail (*Coturnix japonica),* helmeted guineafowl (*Numida meleagris*) and zebra finch (*Taeniopygia guttata).* From the 16,127 protein-coding genes in the Mgal_WU_HG_1.0 gene set, 98% were found to be in an orthogroup. This was the highest percentage of any of the species tested **(Table 4).** Of the 15,417 orthogroups found, 91% include Mgal_WU_HG_1.0 genes. There are also 10 orthogroups that contain only Mgal_WU_HG_1.0 genes, of which two have homologs in the nr database (*MANBAL*, and *POL3*) **(Supplementary File** 1: **Table S2).**

**Table 4:**
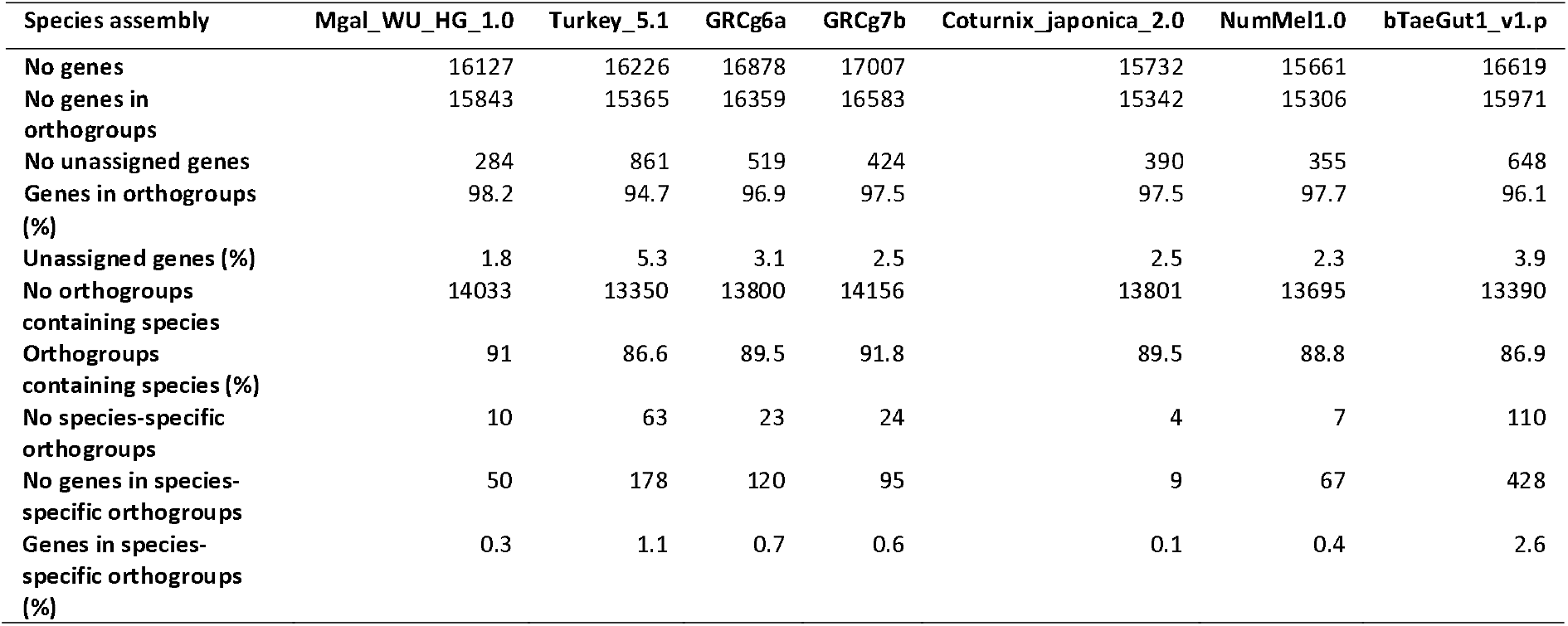
Number of orthogroups found and proportion of genes assigned to each orthogroup per species. Species included: turkey (Mgal_WU_HG_1.0, Turkey_5.1), chicken (GRCg6a, GRCg7b), Japanese quail (Coturnix_japonica_2.0), helmeted guineafowl (NumMel1.0), and zebra finch (bTaeGut1_v1.p)

**Table 5:** Structural variation between the two parental haplotype assemblies. The parent 1 assembly was used as reference and the parent 2 assembly used as the query. Copygain: Copy gain in the query genome, copyloss: copy loss in the query genome.

| Variation type | Count | Length Parent1 | Length Parent2 |
| --- | --- | --- | --- |
| Syntenic regions | 85 | 990,480,776 | 989,217,672 |
| Inversions | 19 | 1,728,932 | 1,525,862 |
| Translocations | 68 | 895,801 | 867,550 |
| Duplications | 397 | 870,354 | 3,179,922 |
| Copy gains | 40 | - | 305,148 |
| Copy losses | 58 | 1,268,056 | - |

#### Gene family contractions and expansions

While most orthogroups studied showed no change in the copy-number of protein coding genes, 29 groups showed expansions or contractions of gene families (27 expansions, 2 contractions) **(Supplementary File 2).** Expanded orthogroups contained proteins involved in important processes in bird development and growth, including gene families involved in cytoskeleton (proteins for feather keratin) (OG0000021, OG0000030, OG0000048), reproduction (involved in spermatogenesis/spermiogenesis) (OG0000005, OG0000883, OG0000065, OG0000973), response to stress (OG0000980), and immunity (OG0000001, OG0000127, OG0001025). Orthogroups OG0000005 and OG0000883 show an expansion of the turkey PHD finger protein 7 (PHF7) gene, which has been shown to be a highly duplicated gene family in the chicken genome [23]. The contracted gene families include one immunoglobin (OG0000001) and one inositol receptor like protein (OG0000070).

### Structural variation between parental haplotypes

The F1 and parental short reads were mapped back to the corresponding assembly with the percentage of mapped reads ranging from 98.73 - 98.91%. Heterozygosity in the F1 assembly was 0.173% (1 heterozygous SNP per 577 bp), while for the parent assemblies lower heterozygosity of 0.117% (parent 1) and 0.107% (parent 2) were found, respectively. This shows that both parental lines generally have low heterozygosity, resulting in a rather low heterozygosity in the F1 as well.

#### Structural variation

The F1 and parent assemblies are completely co-linear **Supplementary File** 1: **Figure S2).** There are no large structural differences (>1 Mbps) between the two parental haplotypes except for a 1.47 Mbp inversion on chromosome 1 (74.28 – 75.74 Mb, **Supplementary File 3)** comprising 25 protein coding genes and 15 lncRNA genes. **Table 3** shows an overview of the number and cumulative length of each type of structural variation.

In total, 231 large structural variations (>10 kb) have been identified between the two parental haplotypes **(Supplementary File 4).** From these, 81 affect the coding sequence of protein coding genes **(Supplementary File 4),** of which 40 have a 1:1 ortholog in chicken. Interestingly, an inversion affecting the coding sequence of the *BLB2* gene (associated with obesity in mice [24]) was found to be homozygous in parent 2 and heterozygous in parent 1. We further identified duplications in the parent 2 haplotype comprising the *TRIM36, GRIA2* and *MAN2B2* gene. Specifically, the parent 2 haplotype exhibits a 20 kb duplication of the 3’ end of *MAN2B2,* a gene which in pigs is associated with ovulation rate [25]. In addition, a 34 Kbp duplication affecting the *GEMIN8* gene in parent 2 was identified. The *GEMIN8* gene product is part of the survival motor neuron (SMN) complex. Moreover, a 53 Kbp duplication was found affecting the 3’ end of the *RIMKLB* gene, resulting in a copy number of 3 in parent 1 but a copy number of >20 in parent 2. In addition, a 100 kb translocation that comprises the *RALYL* gene was identified. The translocated region is found at around 68.2 Mbp on chromosome 5 in parent 1, while it is found at a position around 90.1 Mbp on the same chromosome in parent 2. Finally, an inversion on chromosome 30 of length 187 kb comprises two protein coding genes and one lncRNA.

A full overview of structural variation between the parental haplotypes is provided in **Supplementary File 4.**

#### Loss of function variation

The most common effect of selection is to alter gene expression, leading to phenotypic changes. However, a small proportion of phenotypic variation is due to impaired gene functioning [26]. We assessed the presence of loss-of-function variation (LoF), specifically stop-gained variants affecting genes in either of the two parental haplotypes **(Supplementary File 5).** In total, 138 stop-gained variants affecting 92 genes between the parent1 and parent2 haplotypes were identified. Genes carrying LoF mutations that are especially noteworthy include the *RYR2* gene, which is affected by four LoF variant in parent 2, likely leading to an impaired RYR2 protein. Mutations in the *RYR2* gene are associated with sudden death syndrome in broiler chickens [27]. A second gene worth highlighting is *LRRC41* which, in the parent 2 haplotype, contains a stop-gained variant. Knockouts of this gene lead to increased lean body mass in mice and hence this gene poses an interesting candidate for selection for body weight in turkey.

### Mapping of SNP-chip markers

SNP-chips are useful to study variation (single nucleotide polymorphisms, SNPs) between individuals and are widely applied in genomic selection. We mapped SNP-chip markers from a 65K SNP array (64,800 SNPs; Illumina, Inc.) to Mgal_WU_HG_1.0 **(Supplementary File** 1: **Table S3)** using a custom SNP mapping pipeline (see methods). We mapped 64,536 (99.4%) of the markers to Mgal_WU_HG_1.0. From these, 1,532 markers that were located on unplaced contigs in Turkey_5.1 are now mapped to specific chromosomes in Mgal_WU_HG_1.0, and 415 markers were placed on the new chromosomes 31-35, indicating a higher completeness. More specifically, we were able to place a significant number of new markers, especially on chromosomes 1 (412), 27 (120), 31 (192), and Z (594).

### Distinct genomic landscapes of turkey micro and macrochromosomes

Avian genomes are known to vary greatly in genomic features, especially between the micro and macrochromosomes. We evaluated the genomic landscape of the turkey chromosomes in terms of repeat content, gene density, and gene expression between macro (>40 Mbp), intermediate (>40 Mbp, <20 Mbp), and micro (<20 Mbp) chromosomes. We found that the repeat content of each repeat class in macro, micro and intermediate chromosomes varied highly along the chromosome **(Supplementary File** 1: **Figures S3-10).** Macrochromosomes are enriched for DNA transposons and LINE elements compared to the intermediate and microchromosomes **(Supplementary File** 1: **Figure S3-S4).** In addition, LINE CR1 elements are especially enriched at the tails of macrochromosomes. Microchromosomes are enriched for low complexity **(Supplementary File** 1: **Figure S5),** simple **(Supplementary File** 1: **Figure S7),** and unknown repeats **(Supplementary File** 1: **Figure S10),** the latter especially at the tails of the chromosomes.

In order to assess whether there was a distinction between the type of genes (e.g. tissue specific or housekeeping) in chromosome types, we analysed RNA-seq datasets from 16 tissues (mapping rates in **Supplementary File** 1: **Table S4).** Microchromosomes showed on average higher gene expression than macro and intermediate chromosomes **(Figure 3A**), as well as having a higher relative abundance of housekeeping genes, defined here as genes expressed in at least 13 out of the 16 studied tissues included in this study **(Figure 3B).**

**Figure 3:**
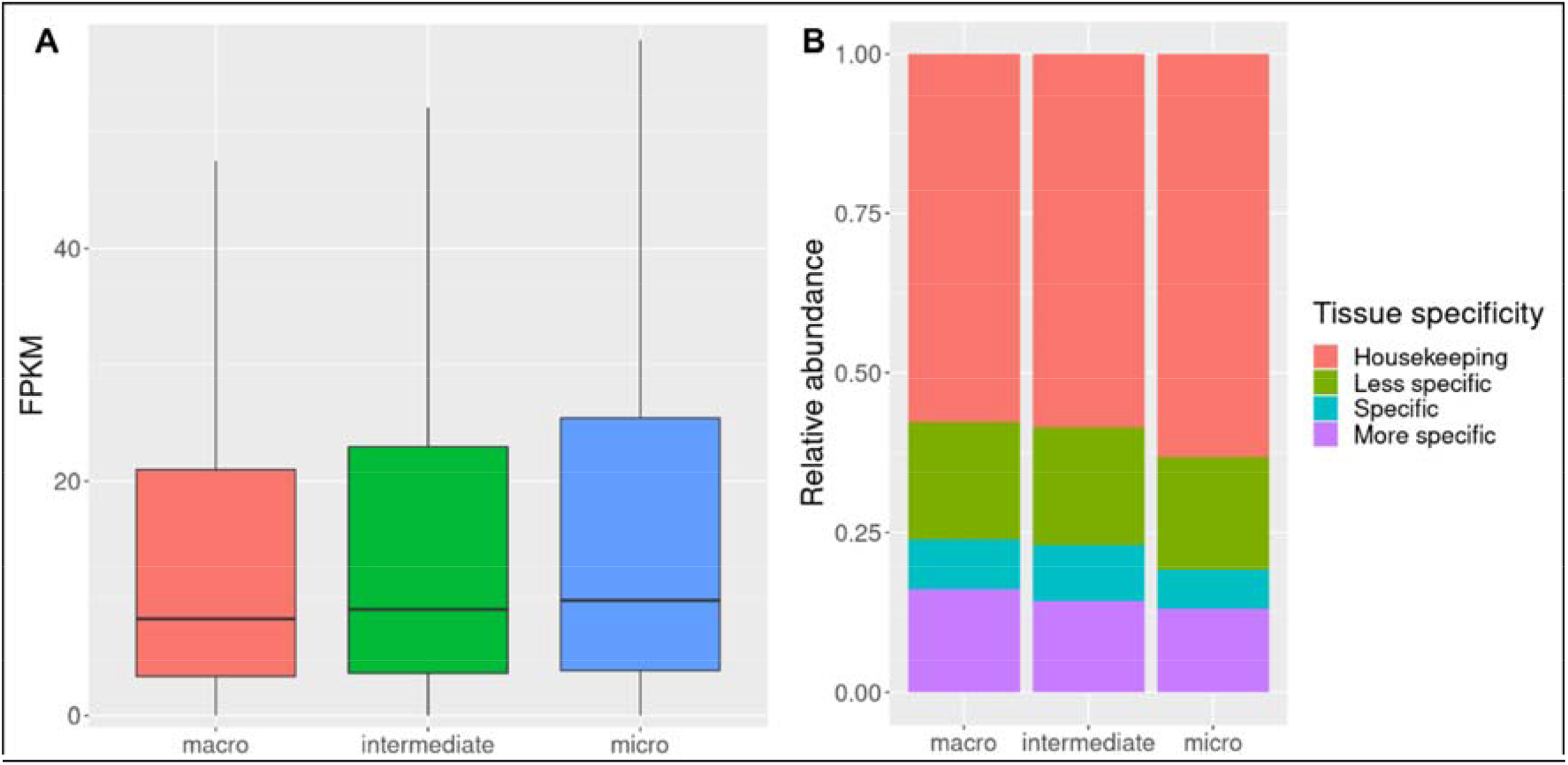
A) Overview of gene expression in macro, intermediate and micro chromosomes. B) Relative abundance of tissue specific genes in each chromosome class. Microchromosomes show higher relative abundance of housekeeping genes when compared with macro and intermediate chromosomes. Number of tissues tested: 16. Housekeeping genes: expressed in at least 13 tissues; less specific genes: expressed in at least 5 tissues and fewer than 13 tissues; specific: expressed in 2 to 5 tissues; more specific: expressed in one or two tissues.

### Conserved synteny within the Galliformes clade

We performed synteny analysis to assess chromosomal and structural rearrangements within a wide range of avian species. Four Galliformes were included: turkey, chicken, Japanese quail, and helmeted guineafowl. Furthermore, two Passeriformes, zebra finch and great tit, and emu, a species from the Casuariiformes order were included. The multi-species synteny plot shows a high degree of synteny between the avian species both on the macro and the microchromosomes, despite the large evolutionary distances **(Figure 4).**

**Figure 4:**
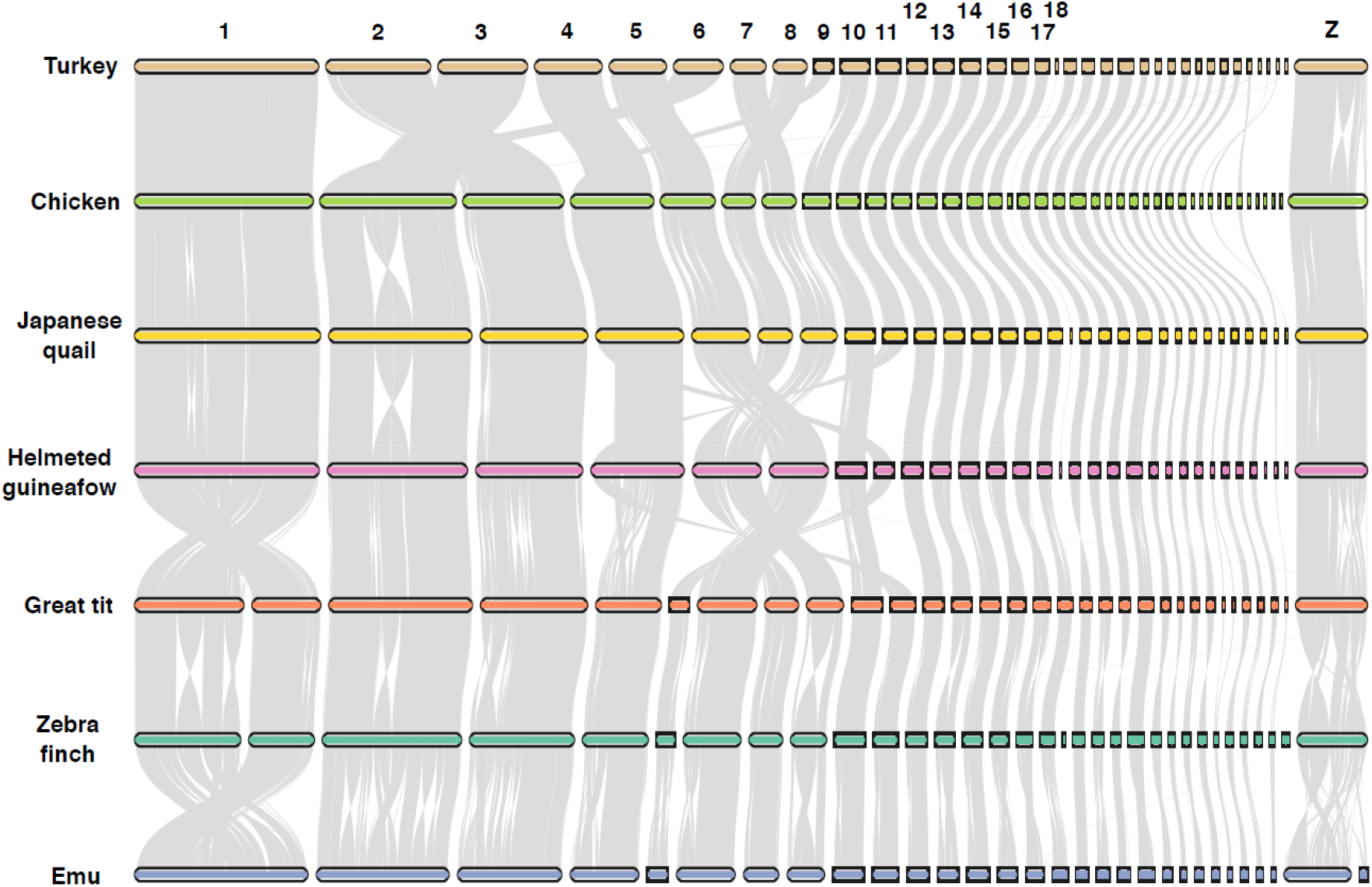
Chromosomal rearrangements across several avian species. Pairwise synteny comparison across 7 birds shows several chromosomal rearrangements. Grey segments represent conserved synteny. Species: turkey (*Meleagris gallopavo),* chicken (*Gallus gallus),* Japanese quail (*Coturnix japonica),* helmeted guineafowl (*Numida meleagris),* great tit (*Parus major),* zebra finch (*Taeniopygia guttata),* and emu (*Dromaius novaehollandiae).*

Of all chromosomes, it is evident that especially the Z chromosome has been prone to large chromosomal rearrangements between avian orders **(Figure 5).** Interestingly, we found a large inversion of around 19 Mbp on the turkey Z chromosome not found in the other Galliformes and songbirds. This is especially striking since rearrangements on the Z chromosome are uncommon within the Galliformes. One region at the tail of the chicken Z chromosome lacks synteny with other Galliformes altogether. This region is enriched in repeat sequences in both chicken and turkey **(Supplementary File 1: Figure S11).**

**Figure 5:**
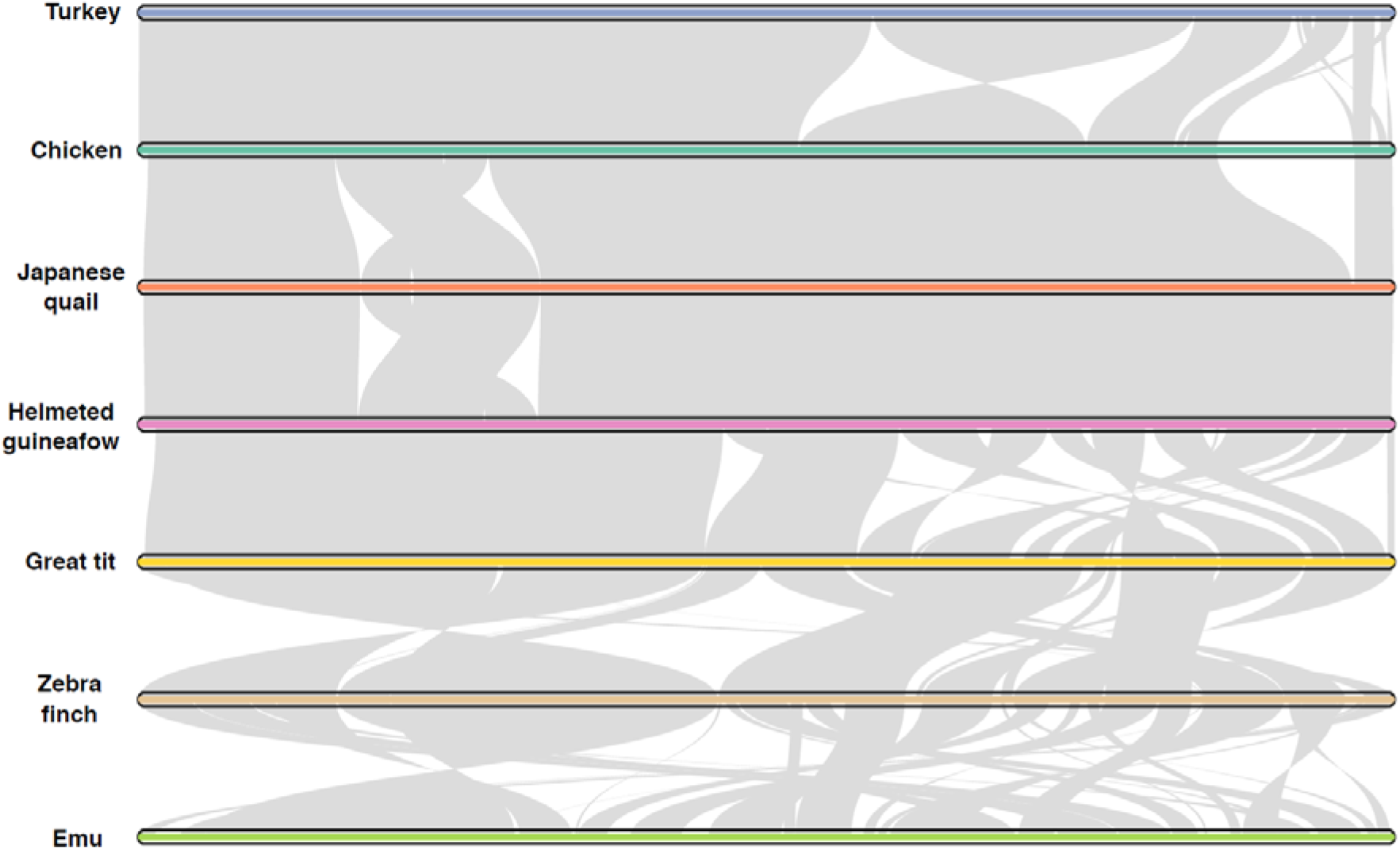
Chromosome Z rearrangements across 7 avian species. Pairwise synteny comparison of the Z chromosome across avian species reveals a large inversion in turkey. Grey segments represent conserved synteny. Species: turkey (*Meleagris gallopavo),* chicken (*Gallus gallus),* Japanese quail (*Coturnix japonica),* helmeted guineafowl (*Numida meleagris),* great tit (*Parus major),* zebra finch (*Taeniopygia guttata),* and emu (*Dromaius novaehollandiae*)

## Discussion

We present a new, chromosome-level, high quality reference assembly for *Meleagris gallopavo,* Mgal_WU_HG_1.0. The trio binning approach has been proven to be a robust method to characterize the two haplotypes of F1 individuals [12]. The quality of the assemblies presented in this study confirms the value of this method in not only providing a quality assembly but also in uncovering structural genomic variation. The Mgal_WU_HG_1.0 assembly is a large improvement over the previous turkey assembly, Turkey_5.1 [5]. The assembly is now comparable in quality and completeness to the chicken reference genome (GRCg6a) and to the recently available GRCg7 genomes. One major limitation of previous turkey assemblies was that they relied on assumptions on high turkey-chicken retained synteny to achieve a chromosome-level assembly. Such assumptions can result in bias, especially when comparing turkey to chicken. Mgal_WU_HG_1.0 does not rely on such comparisons.

Combining long reads and genome-wide chromatin interaction data (Hi-C) enables the capture of chromosome arms in a single contig, resulting in a highly continuous and contiguous chromosome-level assembly. Furthermore, long reads can span long repetitive regions including DNA transposons and LINE elements, as well as large structural variants. Centromeres, however, are too long to traverse reliably in most cases. Thanks to these recent sequencing technologies, we are able to correct a number of wrongly oriented contigs in Turkey_5.1, a phenomenon often observed in short-read based assemblies. The improvements in genome quality, completeness and continuity allow for a more thorough annotation of repeats and gene models. The increase in complete BUSCO genes in Mgal_WU_HG_1.0, compared to Turkey_5.1, indicates a much-improved gene space in the current genome assembly, comparable to the latest chicken genome builds.

Improving genome assemblies improves all analyses that depend on them. One of the reasons to improve the turkey assembly was to better map SNP-chip markers to the genome. SNP-chips are widely used in genomic selection and a better genome representation and gene annotation directly impacts its use for breeding. Specifically, the new turkey genome build overcomes the lack of SNPs mapped to gene-dense microchromosomes, as 85.3% of the SNP markers previously mapped to unplaced scaffolds on Turkey_5.1 are now mapped to chromosomes on Mgal_WU_HG_1.0, especially improving the representation of microchromosomes 31 to 35.

Turkey breeding is done on pure elite lines which can be selected for different purposes. In our study, one parent was from a female breeding line, with more focus on egg production and conformation, whereas the other parent was from a male breeding line focussing on growth and production traits. In producing a commercial product, lines are crossed to produce hybrid offspring that shows the benefit of the breeding goals of both parental lines. In addition, the hybrid offspring benefits from hybrid vigour, resulting from two relatively differentiated lines. For the trio-binning method, having parents that are genetically distinct helps in resolving the haplotypes. Nevertheless, in this study, we present two high quality parental haplotype assemblies where the low heterozygosity of the parents presented no obstacle to resolving the parental haplotypes.

Among the remaining challenges in variation analysis is the characterization of structural variants. The challenge is two-fold. First, these large-scale variants are often not robustly detected using short-read sequencing. Second, individuals usually have sequence that is population specific, and which may not be present in a reference assembly. This can make such large insertions hard to characterize, even by re-sequencing. In the process of assembling Mgal_WU_HG_1.0 we now have reference assemblies for two distinct breeding lines, which should greatly aid in variation analysis. Even though such large structural variants appear to be uncommon between breeding lines, we demonstrate how genes potentially important in breeding may be affected. These genes can be further prioritized in routine genomic breeding practice.

As more genomes are characterized with high accuracy and at a chromosome-level, comparative genomics is increasingly used to study the function of genes and variants, including copy number variants. The new Mgal_WU_HG_1.0 genome assembly was applied to identify orthogroups that have expanded or contracted in turkey compared to other avian species. Expanded orthogroups included various distinct keratin families, encoding major structural proteins of feathers and claws [28]. One gene family comprising the PHD Finger Protein 7 (PHF7) was significantly expanded in turkey. *PHF7* acts during spermiogenesis for histone-to-histone protamine exchange and is a determinant of male fertility in Drosophila and mouse [29], and highly expressed in rooster testis [30]. This gene family was found to be expanded in chicken as well, with distinct gene clusters on five chromosomes [23]. In addition, genes related to immunity and response to stress are expanded in turkey. Further research is needed to disentangle the exact function of these complex gene families.

A characteristic of avian genomes is that they comprise a huge range of chromosome sizes. Interestingly, bird genome organization may be ancestral to all vertebrates [31]. Among the peculiar outcomes is a wide range in e.g. recombination rates, GC-bias, gene densities and variation density throughout the genome [32]. The distinct nature of these features is particularly difficult to study in microchromosomes as they have proven so difficult to characterize. The Mgal_WU_HG_1.0 assembly though, has a better representation of the microchromosomes, allowing a better understanding of functional aspects of genes and other genome elements. We have shown that the microchromosomes have a unique repeat landscape enriched for low complexity, simple, and unknown repeats, especially at the tails of the chromosomes. Together these efforts provide new insights in microchromosome composition and evolution.

Bird genomes have very high retained synteny [33]. This pattern was confirmed in our analysis of the conserved synteny between several Galliformes (turkey, chicken, Japanese quail, helmeted guineafowl) and three outgroups (zebra finch, great tit, emu). Despite the long divergence time that separates turkey and chicken [2], both species have relatively similar karyotypes confirmed by the high structural continuity and relatively little rearrangements between the two birds, even in the microchromosomes. The latter is noteworthy because of the very high recombination rates generally observed in microchromosomes [34], which would suggest that a higher rate of chromosomal rearrangements might be expected but is not observed. Expanding observations to other Galliformes suggest similar degrees of conserved synteny, although comparisons for micro-chromosomes are less accurate due to the more incomplete assembly of these other Galliform species

The Z chromosome presents a moderate yet striking deviation from the observed evolutionary stability. This chromosome exhibits a few rearrangements within the Galliformes and, in line with the findings of Zhang et al. (2011), we observed and validated a large inversion in the turkey Z chromosome. As with the Mgal_WU_HG_1.0 assembly the exact breakpoints of this 19 Mbp inversion on the Z chromosome can now be pinpointed. This inversion is unique for the turkey lineage, and not found in any of the other Galliformes.

In conclusion, the new turkey genome here presented (Mgal_WU_HG_1.0) (and the two parental haplotype assemblies) represents a substantial improvement over the previous assembly and is an important resource with many applications in research and in the turkey breeding industry.

## Methods

### Data and Assembly

To create a high-quality chromosome level genome assembly of *Meleagris gallopavo,* three individuals were sequenced using the trio binning approach - two parents and one F1. The two parents come from two distinct commercial lines from Hendrix Genetics, one male line (parent1) and one female line (parent2). The F1 turkey was sequenced by Dovetail Genomics using PacBio singlemolecule real-time (SMRT) sequencing technology (PacBio Sequel System, RRID:SCR_017989) with a total depth of 270X. We generated short read sequencing data from the F1 (90.4X coverage) and both parents (35.4X, and 39.7X coverage) on an Illumina HiSeq 4000 (HiSeq 4000 System, RRID:SCR_016386). In addition, Hi-C data was generated with a coverage of 32X. An initial assembly was created by Dovetail Genomics using wtdgb2 (WTDBG, RRID:SCR_017225) [13] and scaffolded using the Dovetail *De Novo* Assembly Process, which uses Chicago^®^ and Dovetail Hi-C proximation ligation methods and the HiRise™ scaffolder as described in [14].

### Polishing

Pilon v1.23 (Pilon, RRID:SCR_014731) [36] was used to polish SNPs and indels based on the short Illumina reads from the F1 (twice with parameters--diploid --mindepth 0.7 --fix bases --changes), and indels with the Illumina reads from parent2 because of the higher coverage compared to parent1 (--fix indels).

### Scaffolding

We scaffolded the F1 assembly received by Dovetail Genomics using the Hi-C reads and the PacBio long reads, both from the F1. The Hi-C reads were mapped to the polished assembly based on the Arima Mapping pipeline [37], using BWA-MEM v0.7.17 (BWA, RRID:SCR_010910) [38] with default parameters. The filter_five_end.pl script was used to filter and keep the 5’-end. After filtering, the reads are sorted and paired using the two_read_bam_combiner.pl script. This results in a sorted, paired-end BAM file that has been filtered by mapping quality (mapping quality filter =10). Picard Tools v2.23.4 (Picard, RRID:SCR_006525) [39] - AddOrReplaceReadGroups and MarkDuplicates was used to add a read group and remove duplicates. The mapped Hi-C reads were used to scaffold the assembly with SALSA v2.2 (SALSA, RRID:SCR_022013) [15], which is a scaffolder that uses long range contact information (Hi-C) with parameters -e “GATC”. Redundans v0.14a [40] was used to scaffold the assembly with the PacBio reads with length >40 Kbp and remove redundant contigs from the final assembly. The parameters -l <long reads> --nogaplosing --noscaffolding were used (-noscaffolding skips short read scaffolding).

### Hi-C validation - mis-assemblies

To validate our F1 assembly and look for mis-assemblies we used Hi-C contact maps. Juicer v1.6 (Juicer, RRID:SCR_017226) [41] was used to generate Hi-C contact maps from the Hi-C reads **(Supplementary File** 1: **Figure S1)** and 3D-DNA v180922, a 3D de novo assembly pipeline (3D de novo assembly, RRID:SCR_017227) [42], to scaffold our assembly. Juicebox v1.11.08 (Juicebox, RRID:SCR_021172) [43] was used to visualize the Hi-C contact map and identify mis-assemblies. Each breakpoint in the macrochromosomes was manually checked with Juicebox and JBrowse 1.16.9 (JBrowse, RRID:SCR_001004) [44] to visualize the PacBio read coverage at the breakpoints.

### Haplotype assemblies using trio-binning

TrioCanu (a module from the Canu assembler, v2.1.1) (Canu, RRID:SCR_015880) [12] was used to bin the parental reads to construct parental haplotype assemblies. TrioCanu was run with the short reads from each parent and the F1 PacBio reads with the following options: -p asm genomesize-1.1g. The corrected reads from TrioCanu were mapped to the Triocanu assembly with Minimap2 v2.17-r941 (Minimap2, RRID:SCR_018550) [45], options -x map-pb. LRScaff v1.1.10 [17] was used to scaffold each parent assembly. For both parents the scaffolding was done with these parameters: min_contig_length = 500, identity = 1, min_overlap_length = 400, max_overhang_length = 500, max_end_length = 500, min_supported_links = 2, iqr_time = 3. Duplicated sequences were removed. RagTag v1.1.1 [46] was used for reference-guided scaffolding of each parental assembly, using the F1 assembly as reference. The scaffold module from RagTag was used with default parameters.

### Completeness

#### BUSCO

BUSCO v4.1.2 (BUSCO, RRID:SCR_015008) [19] was run to assess the completeness of the assembly in terms of gene space. BUSCO was run in the genome mode (-m genome) and with the vertebrae (vertebrata_odb10) and aves (aves_odb10) datasets (using the flag -1 <dataset>).

#### Genome comparison - alignment

Genome assembly alignments were generated using D-GENIES v1.3.0 (D-GENIES, RRID:SCR_018967) [47], using minimap2 as the aligner. The chromosomes were sorted on length, and noise (short repeat alignments) was removed from the alignment plot.

#### Structural variation (parents)

Structural variation between the two parental haplotypes was discovered using SyRI v1.5.4 [48]. First, we aligned the two haplotype assemblies using minimap2 with settings -ax asm5 -eqx. Next, we used SyRI to identify structural variation using the minimap2 alignment. Results were plotted using plotsr tool v0.5.3 [49]. Large structural variants were manually validated in JBrowse 1.16.9 [44].

#### Remapping and variant calling

The short Illumina reads from the F1 individual were mapped back to the assembly using BWA-MEM v0.7.17 (BWA, RRID:SCR_010910) [38]. Samblaster v0.1.26 (SAMBLASTER, RRID:SCR_000468) [50] was used to mark duplicates and Samtools v1.14 (SAMTOOLS, RRID:SCR_002105) [51] to sort and index the BAM files. Freebayes v1.3.1 (FreeBayes, RRID:SCR_010761) [52] was used for variant calling with: --use-best-n-alleles 4 --min-base-quality 10 --min-alternate-fraction 0.2 --haplotype-length 0 --ploidy 2 --min-alternate-count 2. The vcffilter module from vcflib v0.00.2019.07.10 [53] was used to discard variants with low phred quality score (<20). Tabix, a module from htslib v1.9 (SAMTOOLS, RRID:SCR_002105) [54] was used to index the VCF files. The stats module from BCFtools v1.9 (SAMtools/BCFtools, RRID:SCR_005227) [55] was used to compute summary statistics of the variant calling. The same process was followed to call variants for each parent. Alignment quality control statistics were computed with QualimMap v.2.2.2-dev (QualiMap, RRID:SCR_001209) [56].

#### SNP-Chip

In order to map SNP markers from the 65K single nucleotide polymorphism (SNP) array (65,000 SNP; Illumina, Inc.) to the new genome build we first aligned the two genome builds (Turkey_5.1 and Mgal_WU_HG_1.0) using nucmer v4.0.0rc1 (MUMmer, RRID:SCR_018171) [57]. Next we converted the delta file to a chain file using mugsy v1.2.3 delta2maf and maf-convert (Mugsy, RRID:SCR_001414) [58]. We used CrossMap v0.6.1 (CrossMap, RRID:SCR_001173) [59] to identify SNP locations on the query Mgal_WU_HG_1.0 assembly. We further performed a blastn v2.11.0+ search (BLASTN, RRID:SCR_001598) [60] to identify the locations of SNPs that could not be mapped from the previous build using the SNPs probe sequences.

#### Annotation and repeats

The genome was annotated with the ENSEMBL annotation pipeline and is available as part of the Ensembl Rapid Release (Ensembl, RRID:SCR_002344) [21]. The transcriptome and proteome evidence used in the annotation are listed in **Supplementary File 6.** We used a custom python script to query the Ensembl rapid release homologue gene page to identify Turkey_5.1 and GRCg6a homologues of all the Mgal_WU_HG_1.0 genes. The BuildDatabase tool from RepeatModeler v1.0.11 (RepeatModeler, RRID:SCR_015027) [20] was used to build a de novo repeat library from our assembly using the Recon and RepeatScout tools. RepeatMasker v4.0.7 (RepeatMasker, RRID:SCR_012954) [61] was used to identify repeats together with the custom build repeat library from RepeatModeler.

#### Orthologues

The proteomes of five bird species were used to infer orthogroups (option -og) using OrthoFinder v2.5.4 (OrthoFinder, RRID:SCR_017118) [62]. The proteomes of the following assemblies were downloaded from Ensembl release 106: turkey -Turkey_5.1; chicken - GRCg6a; Japanese quail - Coturnix_japonica_2.0; helmeted guineafowl - NumMel1.0; zebra finch - bTaeGut1_v1.p. The proteomes for Mgal_WU_HG_1.0 (turkey) and GRCg7b (chicken) were downloaded from the Ensembl rapid release (March 2022). For each orthogroup, the protein isoform with the best alignment based on species similarity, score and expect value was chosen. Turkey-specific orthogroups were analysed by running BLASTp v2.11.0+ (BLASTP, RRID:SCR_001010) [60] against the NR database to identify homologous genes from a wider range of species.

#### Gene family contractions and expansions of protein-coding gene families

Expansions and contractions of protein-coding gene families were assessed by filtering the OrthoFinder results. If the number of proteins for those species is significantly higher than the number of turkey proteins, we consider a contraction in turkey. On the other hand, if the number of proteins in the non-turkey species is significantly lower than the number of turkey genes we consider an expansion in turkey.

#### Distinct genomic landscapes of turkey micro- and macrochromosomes

To better understand the differences between macro (>40 Mbp), intermediate (>40 Mbp, <20 Mbp), and micro (<20 Mbp) chromosomes, we investigated repeat content, gene structure and gene expression.

#### Repeats

A custom repeat library created with RepeatModeler and custom R scripts were used to investigate the differences in repeat content between macro, intermediate and microchromosomes. Each chromosome was split into bins (each bin corresponding to 2% of the chromosome length), allowing us to compare the chromosomes by relative length. We calculated the average repeat content in each bin. An ideogram of the density of each repeat feature was created for macro, intermediate and microchromosomes with the R v4.0.2 (R Project for Statistical Computing, RRID:SCR_001905) [63] package RIdeogram v0.2.2 [64]. RIdeogram calculates feature density in sliding windows (100 Kbp for macro and intermediate chromosomes, 50 Kbp for microchromosomes).

#### Tissue specificity

Expression data for 16 turkey tissues (jejunum, proventriculus, thigh, testis, ileum, pancreas, spleen, breast, brain, heart, thymus, liver, gizzard, duodenum, caecal tonsil, bursa) from a male individual at three developmental stages (14, 21, 28 days post hatch) was downloaded from Bioproject PRJNA259229. Not all tissues were available at all stages: testis was not available at day 21 and caecal tonsil at day 28. HISAT2 v2.2.1 (HISAT2, RRID:SCR_015530) [65] was used to index the assembly (hisat2-build), and align the RNA-seq reads to the assembly. Stringtie v2.1.7 (StringTie, <u>RRID:SCR_016323</u>) [66] was used to assemble transcripts using the aligned reads and Ensembl gene annotation with options -A and -B. A non-redundant set of transcripts was generated with Stringtie’s merge option (--merge), which creates a unified set of transcripts from several samples. Stringtie was run once more, now using this new set of transcripts as the reference annotation file. The resulting table containing the gene abundance of all genes was used in our analysis. We analysed the results through custom R (v4.0.2) scripts. We started by filtering the gene abundance table to keep only the genes that are expressed (FPKM >1). Then we classified genes into housekeeping (expressed in at least 13 tissues), less specific (expressed in at least 5 and in fewer than 13 tissues), specific (expressed in 2 to 5 tissues), and more specific genes (expressed in one or two tissues). The relative abundance of housekeeping/specific genes was calculated by counting the number of genes in these categories in macro, intermediate and microchromosomes and dividing that by the total amount of genes in each chromosome type.

#### Gene structure

We used RIdeogram v0.2.2 [64] and R (v 4.0.2) to compare the gene density between the chromosome classes. RIdeogram calculates gene density in sliding windows, 100 Kbp for macro and intermediate chromosomes, 50 Kbp for microchromosomes.

#### Synteny

The MCscan python pipeline from the JCVI utility libraries v1.1.11 (MCScan, RRID:SCR_017650) [67] was used study chromosomal rearrangements between several bird species: Turkey (*Meleagris gallopavo*), chicken (*Gallus gallus*), Japanese quail (*Coturnix japonica*), helmeted guineafowl (*Numida meleagris*), great tit (*Parus major*), zebra finch (*Taeniopygia guttata*), and emu (*Dromaius novaehollandiae*).

The genome (fasta coding DNA sequence, CDS) and annotation files for these species were obtained from Ensembl release 106. The files for Mgal_WU_HG_1.0 and GRCg7b were obtained from the Ensembl rapid release (April 2022). The annotation file for the emu assembly ZJU1.0 was shared with us from [68]. This annotation file, in combination with the FASTA file obtained from NCBI was used to create the CDS fasta file necessary for the pipeline.

We started by trimming the accession IDs in the FASTA file and converting the GFF3 annotation file to BED format. The jcvi.compara.catalog ortholog and jcvi.compara.synteny screen (with parameters --simple) were used to create the necessary input files for plotting. The synteny plots were created with jcvi.graphics.karyotype using parameter --basepair. To validate the chromosome Z inversion, first, we manually checked the inversion breakpoints (reads spanning) using JBrowse 1.16.9.

## Data Availability

The genome assemblies and sequencing data have been deposited in ENA under Bioproject accession PRJEB42643. The turkey genome and annotations are available through ENSEML Rapid Release (https://rapid.ensembl.org/Meleagris_gallopavo_GCA_905368555.1/).

## Supplementary Files

*Supplementary File 1: Table S1:* Protein homology between Mgal_WU_HG_1.0, Turkey_5.1 and chicken (GRCg6a).

*Supplementary File 1: Table S2:* Blast results of proteins in Mgal_WU_HG_1.0 specific orthogroups.

*Supplementary File 1: Table S3:* Mapping of 65K markers on Mgal5.1 and Mgal_WU_HG_1.0.

*Supplementary File 1: Table S4:* Mapping rate of RNA-seq datasets from 16 tissues to Mgal_WU_HG_1.0. Tissues (jejunum, proventriculus, thigh, testis, ileum, pancreas, spleen, breast, brain, heart, thymus, liver, gizzard, duodenum, caecal tonsil, bursa) are from a male individual at three developmental stages (14, 21, 28 days post hatch).

*Supplementary File 1: Figure S1:* Hi-C contact map of the Mgal_WU_HG_1.0 assembly.

*Supplementary File 1: Figure S2:* Parent 1 vs. parent 2 alignment

*Supplementary File 1: Figure S3:* Average DNA repeat content along the chromosomes for macro, intermediate and microcromosomes.

*Supplementary File 1: Figure S4:* Average LINE repeat content along the chromosomes for macro, intermediate and microcromosomes.

*Supplementary File 1: Figure S5:* Average low complexity repeat content along the chromosomes for macro, intermediate and microcromosomes.

*Supplementary File 1: Figure S6:* Average LTR repeat content along the chromosomes for macro, intermediate and microcromosomes.

*Supplementary File 1: Figure S7:* Average simple repeat content along the chromosomes for macro, intermediate and microcromosomes.

*Supplementary File 1: Figure S8:* Average SINE repeat content along the chromosomes for macro, intermediate and microcromosomes.

*Supplementary File 1: Figure S9:* Average snRNA repeat content along the chromosomes for macro, intermediate and microcromosomes.

*Supplementary File 1: Figure S10:* Average unknown repeat content along the chromosomes for macro, intermediate and microcromosomes.

*Supplementary File 1: Figure S11:* Schematic view of Gal7b chromosome Z and representation of several biotypes of genes and genomic features (Ensembl, rapid release 15^th^ June 2022, accessed on 27^th^ June 2022).

*Supplementary File 2:* Gene family expansions and contractions.

*Supplementary File 3:* Syri output showing structural variation between two parent haplotypes.

*Supplementary File 4:* Structural variation table between parent haplotypes.

*Supplementary File 5:* Stop-gained variants identified in either or one of the two parent assemblies.

*Supplementary File 6:* Transcriptome and proteome evidence used for ENSEMBL Annotation.

## Competing Interests

J. Mohr and B.J. Wood were employed by Hybrid Turkeys and M.C.A.M Bink was employed by Hendrix Genetics Research. Both institutes are part of one of the funders (Hendrix Genetics). All authors declare that the results are presented in full and as such present no conflict of interest. The other Breed4Food partners Cobb Europe, CRV, Topigs Norsvin, declare to have no competing interests for this study.

## Funding

This research was funded by the STW-Breed4Food Partnership, project number 14283: From sequence to phenotype: detecting deleterious variation by prediction of functionality. This study was financially supported by NWO-TTW and the Breed4Food partners Cobb Europe, CRV, Hendrix Genetics and Topigs Norsvin.

## Ethical Statement

Ethical review and approval were not required for sample collection since the data used in this study has been obtained as part of routine data collection from Hybrid Turkeys’ breeding programmes, and not specifically for the purpose of this project. Therefore, approval of an ethics committee was not mandatory.

## Authors’ Contributions

MAMG designed, coordinated, and managed the project; JM and BJW were involved in data collection and preparation; RPMAC was involved in data collection and wet lab work; HJM provided valuable input regarding the analyses and manuscript; CPB and MFLD performed the analysis and drafted the manuscript. All authors read and approved the final manuscript.

## Acknowledgements

We are grateful to Luohao Xu (Key Laboratory of Freshwater Fish Reproduction and Development, Southwest University, Chongqing) for sharing the annotation file for ZJU1.0. We thank the ENSEMBL support team for providing details on the annotation.

## Abbreviations

BED: Browser Extensible Data;
BLAST: Basic Local Alignment Search Tool;
BLASTN: BLAST search of nucleotide database(s);
BLASTP: BLAST search protein databases using a protein query;
bp: base pairs;
BUSCO: Benchmarking Universal Single-Copy Orthologs;
BWA: Burrows-Wheeler Aligner;
CDS: coding sequence;
F1: Filial 1, first offspring from a cross;
FPKM: Fragments Per Kilobase Million;
GC: guanine-cytosine;
GFF3: general feature format, version 3;
GWAS: genome wide association studies;
Hi-C: chromosome conformation capture;
INDEL: insertion or deletion;
Kbp: kilo base pairs;
LINE: Long interspersed nuclear elements;
InRNA: long non-coding RNA;
LoF: loss of function;
Mbp: megabase pairs;
NCBI: National Center for Biotechnology Information;
PacBio: Pacific Biosciences;
SMN: survival motor neuron;
SMRT: single molecule real time;
SNP: single nucleotide polymorphism;
VCF: variant call format.

