## Supplementary material for "A new haplotype-resolved turkey genome to enable turkey genetics and genomics research": SupplementaryFile1.docx

Table S1: Protein homology between Mgal_WU_HG_1.0, Turkey_5.1 and chicken (GRCg6a).

| Homologues in: | No genes |
| --- | --- |
| Turkey_5.1 | 13281 |
| GRCg6a | 13915 |
| Turkey_5.1 and GRCg6a | 12372 |
| Total in dataset | 16127 |

Table S2: Blast results of proteins in Mgal_WU_HG_1.0 specific orthogroups.

| **Mgal_WU_HG_1.0 specific orthogroups** | **N genes** | **Best hit** | **Species** | **Score** | **expect** |  |
| --- | --- | --- | --- | --- | --- | --- |
| OG0000987 | 11 | E3 ubiquitin-protein ligase Topors-like | Turkey | 46.6 bits (109) | 0.001 |  |
| OG0011308 | 7 | CCNF (cyclin F) | Marbled wood quail | 63.9 bits (154) | 2e-09 |  |
| OG0011309* | 7 | Probable E3 ubiquitin-protein ligase HERC4 isoform X3 | Northern white-cheeked gibbon | 59.3 bits (142) | 2e-09 |  |
| **OG0013797** | **5** | **Protein MANBAL-like** | **Turkey** | **110 bits (275)** | **2e-29** |  |
| OG0013798* | 5 | Wingless-type MMTV integration site family, member 7B | Homo sapiens | 60.5 bits (145) | 1e-8 |  |
| OG0013799* | 5 | hypothetical protein PVAP13_4KG009300 | Switchgrass | 48.5 bits (114) | 5e-04 |  |
| OG0014231* | 4 | hCG1816008 | Homo sapiens | 51.2 bits (121) | 1e-06 |  |
| **OG0015325** | **2** | **POL3 protein / POL2 protein** | **White-crested guan/Pallas's sandgrouse** | **251 bits (642)** | **8e-83** | Catalytic component of DNA polymerase delta (DNA polymerase III) which participates in chromosomal DNA replication |
| OG0015337* | 2 | Hypothetical protein EGK_13122 | Rhesus macaque | 56.6 bits (135) | 2e-08 |  |
| OG0015340 | 2 | Hypothetical protein AXJ14_gp023 | Geobacillus virus E3 | 35.8 bits (81) | 5.6 |  |

*****	No significant matches within Aves

Table S3: Mapping of 65K markers on Mgal5.1 and Mgal_WU_HG_1.0.

| **Chr** | **# SNPs on Mgal5.1** | **#SNPs lost in Mgal_WU_HG_1.0** | **# SNPs gained in Mgal_WU_HG_1.0** | **# SNPs on Mgal_WU_HG_1.0** |
| --- | --- | --- | --- | --- |
| 1 | 7068 | 44 | 412 | 7436 |
| 2 | 4085 | 44 | 125 | 4166 |
| 3 | 3468 | 48 | 122 | 3542 |
| 4 | 2450 | 41 | 37 | 2446 |
| 5 | 2143 | 13 | 32 | 2162 |
| 6 | 3758 | 16 | 54 | 3796 |
| 7 | 2678 | 20 | 2 | 2660 |
| 8 | 2393 | 11 | 20 | 2402 |
| 9 | 1231 | 21 | 10 | 1220 |
| 10 | 2123 | 17 | 18 | 2124 |
| 11 | 2732 | 57 | 6 | 2681 |
| 12 | 2179 | 6 | 1 | 2174 |
| 13 | 2652 | 11 | 6 | 2647 |
| 14 | 2048 | 8 | 3 | 2043 |
| 15 | 2417 | 15 | 4 | 2406 |
| 16 | 2043 | 14 | 1 | 2030 |
| 17 | 1905 | 55 | 3 | 1853 |
| 18 | 41 | 5 | 3 | 39 |
| 19 | 1400 | 9 | 5 | 1396 |
| 20 | 1416 | 11 | 5 | 1410 |
| 21 | 1447 | 9 | 9 | 1447 |
| 22 | 1629 | 9 | 4 | 1624 |
| 23 | 1013 | 3 | 3 | 1013 |
| 24 | 680 | 5 | 41 | 716 |
| 25 | 798 | 9 | 2 | 791 |
| 26 | 1004 | 5 | 2 | 1001 |
| 27 | 175 | 2 | 120 | 293 |
| 28 | 721 | 4 | 15 | 732 |
| 29 | 692 | 24 | 50 | 718 |
| 30 | 660 | 35 | 14 | 639 |
| 31 | 0 | 0 | 192 | 192 |
| 32 | 0 | 0 | 53 | 53 |
| 33 | 0 | 0 | 82 | 82 |
| 34 | 0 | 0 | 59 | 59 |
| 35 | 0 | 0 | 29 | 29 |
| Z | 3954 | 34 | 594 | 4514 |
| Unplaced | 1796 | 1532 | 682 | 264 |

Table S4: Mapping rate of RNA-seq datasets from 16 tissues to Mgal_WU_HG_1.0. Tissues (jejunum, proventriculus, thigh, testis, ileum, pancreas, spleen, breast, brain, heart, thymus, liver, gizzard, duodenum, caecal tonsil, bursa ) are from a male individual at three developmental stages (14, 21, 28 days post hatch).

| Sample ID | Sample information | Total reads | Unmapped | Aligned one time | Aligned multiple | Alignment rate |
| --- | --- | --- | --- | --- | --- | --- |
| SRR1570211 | Gizzard_D14_M | 23328241 | 2234476 | 20808672 | 285093 | 90.42 |
| SRR1570212 | Gizzard_D14_M | 18116364 | 2686852 | 15214616 | 214896 | 85.17 |
| SRR1570213 | Gizzard_D14_M | 10899753 | 1386769 | 8958800 | 554184 | 87.28 |
| SRR1570214 | Gizzard_D14_M | 18071777 | 1525937 | 16293973 | 251867 | 91.56 |
| SRR1570219 | Gizzard_D28_M | 717 | 106 | 598 | 13 | 85.22 |
| SRR1570220 | Gizzard_D28_M | 23507958 | 2783996 | 20389772 | 334190 | 88.16 |
| SRR1570221 | Gizzard_D28_M | 18162922 | 3618220 | 14251886 | 292816 | 80.08 |
| SRR1570222 | Gizzard_D28_M | 13396354 | 1502254 | 11692480 | 201620 | 88.79 |
| SRR1570243 | Thymus_D14_M | 10611043 | 416129 | 10010887 | 184027 | 96.08 |
| SRR1570244 | Thymus_D14_M | 13886003 | 569368 | 13106151 | 210484 | 95.9 |
| SRR1570245 | Thymus_D14_M | 14159584 | 777577 | 13174026 | 207981 | 94.51 |
| SRR1570246 | Thymus_D14_M | 16276374 | 651425 | 15224105 | 400844 | 96 |
| SRR1570250 | Thymus_D28_M | 14738153 | 859306 | 13651411 | 227436 | 94.17 |
| SRR1570251 | Thymus_D28_M | 8304357 | 372628 | 7804905 | 126824 | 95.51 |
| SRR1570272 | Thigh_D14_M | 11067296 | 1863170 | 8385257 | 818869 | 83.17 |
| SRR1570273 | Thigh_D14_M | 11440354 | 1592106 | 8547853 | 1300395 | 86.08 |
| SRR1570274 | Thigh_D14_M | 15900 | 2583 | 11608 | 1709 | 83.75 |
| SRR1570275 | Thigh_D14_M | 15889040 | 2384107 | 11896631 | 1608302 | 85 |
| SRR1570280 | Thigh_D28_M | 8841208 | 1641948 | 6861257 | 338003 | 81.43 |
| SRR1570281 | Thigh_D28_M | 14738588 | 2117385 | 10842910 | 1778293 | 85.63 |
| SRR1570282 | Thigh_D28_M | 11676185 | 1634220 | 8889302 | 1152663 | 86 |
| SRR1570283 | Thigh_D28_M | 15389854 | 2610416 | 11430348 | 1349090 | 83.04 |
| SRR1570284 | Testies_D14_M | 10291093 | 456132 | 9062982 | 771979 | 95.57 |
| SRR1570285 | Testies_D14_M | 16612360 | 806948 | 14561413 | 1243999 | 95.14 |
| SRR1570286 | Testies_D14_M | 18648918 | 938619 | 16095778 | 1614521 | 94.97 |
| SRR1570287 | Testies_D14_M | 15222120 | 757928 | 13410649 | 1053543 | 95.02 |
| SRR1570288 | Testies_D28_M | 17417908 | 820448 | 15459582 | 1137878 | 95.29 |
| SRR1570289 | Testies_D28_M | 12551709 | 619990 | 11105642 | 826077 | 95.06 |
| SRR1570290 | Testies_D28_M | 10954499 | 657102 | 9818545 | 478852 | 94 |
| SRR1570291 | Testies_D28_M | 11176553 | 420155 | 10080114 | 676284 | 96.24 |
| SRR1570312 | Proventriculus_D14_M | 8105954 | 380248 | 7204070 | 521636 | 95.31 |
| SRR1570313 | Proventriculus_D14_M | 9639376 | 1311711 | 7471312 | 856353 | 86.39 |
| SRR1570314 | Proventriculus_D14_M | 13904711 | 2231074 | 10607168 | 1066469 | 83.95 |
| SRR1570315 | Proventriculus_D14_M | 13616829 | 1381221 | 10914618 | 1320990 | 89.86 |
| SRR1570320 | Proventriculus_D28_M | 7126648 | 621878 | 5692213 | 812557 | 91.27 |
| SRR1570321 | Proventriculus_D28_M | 11549691 | 1722063 | 8823933 | 1003695 | 85.09 |
| SRR1570322 | Proventriculus_D28_M | 9965447 | 922162 | 7895811 | 1147474 | 90.75 |
| SRR1570323 | Proventriculus_D28_M | 12625687 | 1218882 | 9948035 | 1458770 | 90.35 |
| SRR1570324 | Proventriculus_D28_M | 12403955 | 1200700 | 9765405 | 1437850 | 90.32 |
| SRR1570345 | Spleen_D14_M | 16333925 | 1111492 | 14961216 | 261217 | 93.2 |
| SRR1570346 | Spleen_D14_M | 11677861 | 786605 | 10702646 | 188610 | 93.26 |
| SRR1570347 | Spleen_D14_M | 11550529 | 745376 | 10621730 | 183423 | 93.55 |
| SRR1570348 | Spleen_D14_M | 14156741 | 947043 | 12084725 | 1124973 | 93.31 |
| SRR1570353 | Spleen_D28_M | 10171696 | 1329324 | 8613366 | 229006 | 86.93 |
| SRR1570354 | Spleen_D28_M | 12616921 | 877116 | 11420809 | 318996 | 93.05 |
| SRR1570355 | Spleen_D28_M | 9668237 | 702020 | 8687793 | 278424 | 92.74 |
| SRR1570376 | Pancreas_D14_M | 17566059 | 1268541 | 11156689 | 5140829 | 92.78 |
| SRR1570377 | Pancreas_D14_M | 20499451 | 1376127 | 13125246 | 5998078 | 93.29 |
| SRR1570378 | Pancreas_D14_M | 17835492 | 1170380 | 11285445 | 5379667 | 93.44 |
| SRR1570379 | Pancreas_D14_M | 22593692 | 1260265 | 14501542 | 6831885 | 94.42 |
| SRR1570384 | Pancreas_D28_M | 17341022 | 910237 | 11001016 | 5429769 | 94.75 |
| SRR1570385 | Pancreas_D28_M | 17155106 | 1209670 | 11934932 | 4010504 | 92.95 |
| SRR1570386 | Pancreas_D28_M | 26155312 | 1591837 | 16297722 | 8265753 | 93.91 |
| SRR1570387 | Pancreas_D28_M | 13217071 | 914113 | 8231127 | 4071831 | 93.08 |
| SRR1570416 | Jejunum_D14_M | 14704077 | 1133087 | 13271434 | 299556 | 92.29 |
| SRR1570417 | Jejunum_D14_M | 21070721 | 3339440 | 17257644 | 473637 | 84.15 |
| SRR1570418 | Jejunum_D14_M | 14991443 | 1306411 | 13319418 | 365614 | 91.29 |
| SRR1570419 | Jejunum_D14_M | 21421194 | 1366889 | 19256822 | 797483 | 93.62 |
| SRR1570424 | Jejunum_D28_M | 19412962 | 1885451 | 17097716 | 429795 | 90.29 |
| SRR1570425 | Jejunum_D28_M | 16745991 | 2301963 | 13910461 | 533567 | 86.25 |
| SRR1570426 | Jejunum_D28_M | 19688505 | 1994163 | 17203910 | 490432 | 89.87 |
| SRR1570427 | Jejunum_D28_M | 26381361 | 2187936 | 23037359 | 1156066 | 91.71 |
| SRR1570448 | Ileum_D14_M | 15339255 | 583691 | 14335954 | 419610 | 96.19 |
| SRR1570449 | Ileum_D14_M | 19774598 | 1374648 | 17890981 | 508969 | 93.05 |
| SRR1570450 | Ileum_D14_M | 12880983 | 868481 | 11759675 | 252827 | 93.26 |
| SRR1570451 | Ileum_D14_M | 17423310 | 1309358 | 15645977 | 467975 | 92.49 |
| SRR1570456 | Ileum_D28_M | 15280824 | 1206563 | 13594433 | 479828 | 92.1 |
| SRR1570457 | Ileum_D28_M | 16085711 | 1807203 | 13973866 | 304642 | 88.77 |
| SRR1570458 | Ileum_D28_M | 17600794 | 1353077 | 15812390 | 435327 | 92.31 |
| SRR1570459 | Ileum_D28_M | 10776435 | 762775 | 9699355 | 314305 | 92.92 |
| SRR1570479 | Heart_D14_M | 18484864 | 5911488 | 12060297 | 513079 | 68.02 |
| SRR1570480 | Heart_D14_M | 22638961 | 6166929 | 15644770 | 827262 | 72.76 |
| SRR1570481 | Heart_D14_M | 17130404 | 5465238 | 11278825 | 386341 | 68.1 |
| SRR1570482 | Heart_D14_M | 14475571 | 5618365 | 8529523 | 327683 | 61.19 |
| SRR1570487 | Heart_D28_M | 15356119 | 4904222 | 10052959 | 398938 | 68.06 |
| SRR1570488 | Heart_D28_M | 17392041 | 5963845 | 10759686 | 668510 | 65.71 |
| SRR1570489 | Heart_D28_M | 18069822 | 3946840 | 12887722 | 1235260 | 78.16 |
| SRR1570509 | Duodenum_D14_M | 25030248 | 3280079 | 21194133 | 556036 | 86.9 |
| SRR1570510 | Duodenum_D14_M | 17888568 | 2593204 | 14870626 | 424738 | 85.5 |
| SRR1570511 | Duodenum_D14_M | 22211069 | 2304579 | 19338228 | 568262 | 89.62 |
| SRR1570512 | Duodenum_D14_M | 27486526 | 3513863 | 23181408 | 791255 | 87.22 |
| SRR1570516 | Duodenum_D28_M | 23819844 | 3175294 | 20057962 | 586588 | 86.67 |
| SRR1570517 | Duodenum_D28_M | 14693978 | 1748896 | 12630782 | 314300 | 88.1 |
| SRR1570518 | Duodenum_D28_M | 9531405 | 2128117 | 7164263 | 239025 | 77.67 |
| SRR1570519 | Duodenum_D28_M | 13079104 | 4875958 | 7794505 | 408641 | 62.72 |
| SRR1570539 | Cecaltonsil_D14_M | 11332863 | 346504 | 10794750 | 191609 | 96.94 |
| SRR1570540 | Cecaltonsil_D14_M | 10877037 | 2118445 | 8556420 | 202172 | 80.52 |
| SRR1570541 | Cecaltonsil_D14_M | 11191194 | 818324 | 10159930 | 212940 | 92.69 |
| SRR1570542 | Cecaltonsil_D14_M | 15017628 | 1216119 | 13485007 | 316502 | 91.9 |
| SRR1570564 | Bursa_D14_M | 11500418 | 973394 | 6994935 | 3532089 | 91.54 |
| SRR1570565 | Bursa_D14_M | 14609285 | 945322 | 12738805 | 925158 | 93.53 |
| SRR1570566 | Bursa_D14_M | 15875406 | 1282244 | 14424983 | 168179 | 91.92 |
| SRR1570567 | Bursa_D14_M | 19308949 | 407448 | 15021488 | 3880013 | 97.89 |
| SRR1570571 | Bursa_D28_M | 14416585 | 653726 | 6947991 | 6814868 | 95.47 |
| SRR1570572 | Bursa_D28_M | 19664715 | 1419200 | 18023354 | 222161 | 92.78 |
| SRR1570593 | Brain_D14_M | 15269278 | 1962359 | 12997128 | 309791 | 87.15 |
| SRR1570594 | Brain_D14_M | 8329351 | 1221333 | 6940949 | 167069 | 85.34 |
| SRR1570595 | Brain_D14_M | 13412813 | 1032836 | 12048509 | 331468 | 92.3 |
| SRR1570600 | Brain_D21_M | 14817236 | 1852543 | 12574139 | 390554 | 87.5 |
| SRR1570601 | Brain_D21_M | 11873796 | 2486871 | 8994456 | 392469 | 79.06 |
| SRR1570602 | Brain_D21_M | 10158280 | 1026149 | 7999945 | 1132186 | 89.9 |
| SRR1570603 | Brain_D21_M | 11744437 | 1564238 | 9936841 | 243358 | 86.68 |
| SRR1570608 | Brain_D28_M | 15973833 | 2784379 | 12801223 | 388231 | 82.57 |
| SRR1570609 | Brain_D28_M | 9987839 | 1525599 | 8299172 | 163068 | 84.73 |
| SRR1570610 | Brain_D28_M | 7989375 | 1075494 | 6706088 | 207793 | 86.54 |
| SRR1570611 | Brain_D28_M | 9486277 | 1462033 | 7778367 | 245877 | 84.59 |
| SRR1570632 | Liver_D14_M | 17769974 | 2318772 | 14927998 | 523204 | 86.95 |
| SRR1570633 | Liver_D14_M | 21123365 | 2923179 | 17649575 | 550611 | 86.16 |
| SRR1570634 | Liver_D14_M | 30567951 | 3602222 | 26265058 | 700671 | 88.22 |
| SRR1570635 | Liver_D14_M | 2616208 | 101662 | 2421888 | 92658 | 96.11 |
| SRR1570640 | Liver_D21_M | 19536192 | 2539960 | 16360833 | 635399 | 87 |
| SRR1570641 | Liver_D21_M | 20350315 | 3066252 | 16779925 | 504138 | 84.93 |
| SRR1570642 | Liver_D21_M | 13524327 | 1548065 | 11557385 | 418877 | 88.55 |
| SRR1570643 | Liver_D21_M | 12398008 | 1599283 | 10530493 | 268232 | 87.1 |
| SRR1570648 | Liver_D28_M | 16727592 | 2127127 | 13992860 | 607605 | 87.28 |
| SRR1570649 | Liver_D28_M | 19984930 | 2424281 | 16983332 | 577317 | 87.87 |
| SRR1570650 | Liver_D28_M | 38345151 | 5100613 | 32261006 | 983532 | 86.7 |
| SRR1570651 | Liver_D28_M | 18992592 | 2101725 | 16257711 | 633156 | 88.93 |
| SRR1570655 | Bursa_D21_M | 14946172 | 1121620 | 13544025 | 280527 | 92.5 |
| SRR1570656 | Bursa_D21_M | 15998205 | 1178033 | 14560230 | 259942 | 92.64 |
| SRR1570657 | Bursa_D21_M | 15924728 | 1250152 | 14418249 | 256327 | 92.15 |
| SRR1570662 | Cecaltonsil_D21_M | 11230735 | 1694509 | 9249767 | 286459 | 84.91 |
| SRR1570663 | Cecaltonsil_D21_M | 23816803 | 2114228 | 19434901 | 2267674 | 91.12 |
| SRR1570664 | Cecaltonsil_D21_M | 12587641 | 2379373 | 9987210 | 221058 | 81.1 |
| SRR1570669 | Duodenum_D21_M | 24752869 | 2697157 | 21455067 | 600645 | 89.1 |
| SRR1570670 | Duodenum_D21_M | 19874160 | 2455661 | 16972372 | 446127 | 87.64 |
| SRR1570671 | Duodenum_D21_M | 16136994 | 2168195 | 13519696 | 449103 | 86.56 |
| SRR1570676 | Gizzard_D21_M | 18722970 | 2236622 | 16273743 | 212605 | 88.05 |
| SRR1570677 | Gizzard_D21_M | 35646337 | 6963092 | 28254097 | 429148 | 80.47 |
| SRR1570678 | Gizzard_D21_M | 12319807 | 2061816 | 10070278 | 187713 | 83.26 |
| SRR1570679 | Gizzard_D21_M | 24944841 | 3909575 | 20674169 | 361097 | 84.33 |
| SRR1570684 | Heart_D21_M | 21086553 | 6969250 | 13551428 | 565875 | 66.95 |
| SRR1570685 | Heart_D21_M | 20450072 | 5222670 | 14568728 | 658674 | 74.46 |
| SRR1570686 | Heart_D21_M | 16172876 | 2563146 | 13081417 | 528313 | 84.15 |
| SRR1570687 | Heart_D21_M | 13381562 | 4116525 | 8891760 | 373277 | 69.24 |
| SRR1570692 | Ileum_D21_M | 16739336 | 1386931 | 15058104 | 294301 | 91.71 |
| SRR1570693 | Ileum_D21_M | 17623899 | 1450681 | 15848254 | 324964 | 91.77 |
| SRR1570694 | Ileum_D21_M | 18413501 | 1449217 | 16620760 | 343524 | 92.13 |
| SRR1570695 | Ileum_D21_M | 15429467 | 1335656 | 13834384 | 259427 | 91.34 |
| SRR1570700 | Jejunum_D21_M | 12573901 | 1072344 | 11177738 | 323819 | 91.47 |
| SRR1570701 | Jejunum_D21_M | 16220020 | 1435092 | 14322270 | 462658 | 91.15 |
| SRR1570702 | Jejunum_D21_M | 16740482 | 1369811 | 14916607 | 454064 | 91.82 |
| SRR1570703 | Jejunum_D21_M | 6697517 | 524194 | 5958999 | 214324 | 92.17 |
| SRR1570725 | Breast_D14_M | 8398350 | 1008640 | 7095218 | 294492 | 87.99 |
| SRR1570726 | Breast_D14_M | 11242282 | 1051823 | 9764034 | 426425 | 90.64 |
| SRR1570727 | Breast_D14_M | 10604446 | 1212365 | 9082105 | 309976 | 88.57 |
| SRR1570728 | Breast_D14_M | 11058025 | 1453627 | 9253025 | 351373 | 86.85 |
| SRR1570733 | Breast_D28_M | 10922458 | 1331535 | 9285913 | 305010 | 87.81 |
| SRR1570734 | Breast_D28_M | 10778225 | 1229415 | 9147354 | 401456 | 88.59 |
| SRR1570735 | Breast_D28_M | 11308550 | 1408348 | 9560154 | 340048 | 87.55 |
| SRR1570736 | Breast_D28_M | 11349746 | 1463991 | 9552223 | 333532 | 87.1 |
| SRR1661425 | Breast_D21_M | 11188693 | 1049761 | 9675323 | 463609 | 90.62 |
| SRR1661426 | Breast_D21_M | 11859847 | 1661955 | 9877255 | 320637 | 85.99 |
| SRR1661427 | Breast_D21_M | 12110962 | 2319775 | 9510160 | 281027 | 80.85 |
| SRR1661428 | Breast_D21_M | 11889516 | 1666043 | 9901052 | 322421 | 85.99 |
| SRR1661433 | Pancreas_D21_M | 17428117 | 1137442 | 11144878 | 5145797 | 93.47 |
| SRR1661434 | Pancreas_D21_M | 14988277 | 873128 | 9836760 | 4278389 | 94.17 |
| SRR1661435 | Pancreas_D21_M | 19066483 | 1394323 | 12196558 | 5475602 | 92.69 |
| SRR1661436 | Pancreas_D21_M | 16883792 | 1087137 | 10824204 | 4972451 | 93.56 |
| SRR1661440 | Proventriculus_D21_M | 10856366 | 2400222 | 6526922 | 1929222 | 77.89 |
| SRR1661441 | Proventriculus_D21_M | 11951154 | 1348025 | 9213772 | 1389357 | 88.72 |
| SRR1661442 | Proventriculus_D21_M | 8496342 | 1030792 | 6626530 | 839020 | 87.87 |
| SRR1661443 | Proventriculus_D21_M | 7611077 | 872185 | 5887181 | 851711 | 88.54 |
| SRR1661447 | Spleen_D21_M | 20932512 | 1702163 | 18438420 | 791929 | 91.87 |
| SRR1661448 | Spleen_D21_M | 12465746 | 1801620 | 10166575 | 497551 | 85.55 |
| SRR1661453 | Thigh_D21_M | 9634946 | 2036072 | 7265832 | 333042 | 78.87 |
| SRR1661454 | Thigh_D21_M | 13366087 | 2349893 | 9731089 | 1285105 | 82.42 |
| SRR1661455 | Thigh_D21_M | 867 | 142 | 646 | 79 | 83.62 |
| SRR1661456 | Thigh_D21_M | 15137603 | 2798014 | 11139170 | 1200419 | 81.52 |
| SRR1661460 | Thymus_D21_M | 15075023 | 910159 | 13886951 | 277913 | 93.96 |
| SRR1661461 | Thymus_D21_M | 12886076 | 777335 | 11905204 | 203537 | 93.97 |
| SRR1661462 | Thymus_D21_M | 13917031 | 976377 | 11936181 | 1004473 | 92.98 |

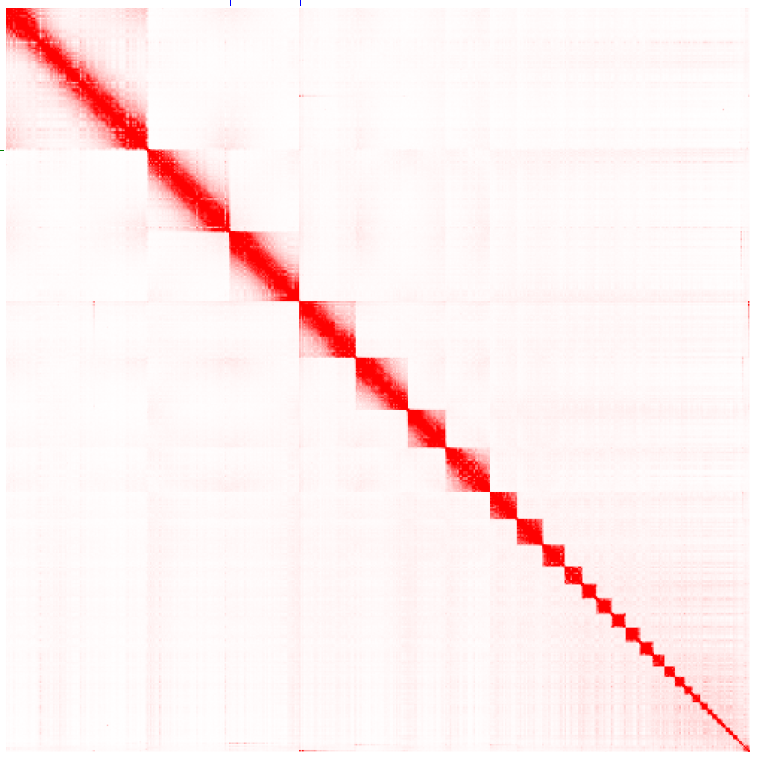

Figure S1: Hi-C contact map of the Mgal_WU_HG_1.0 assembly.

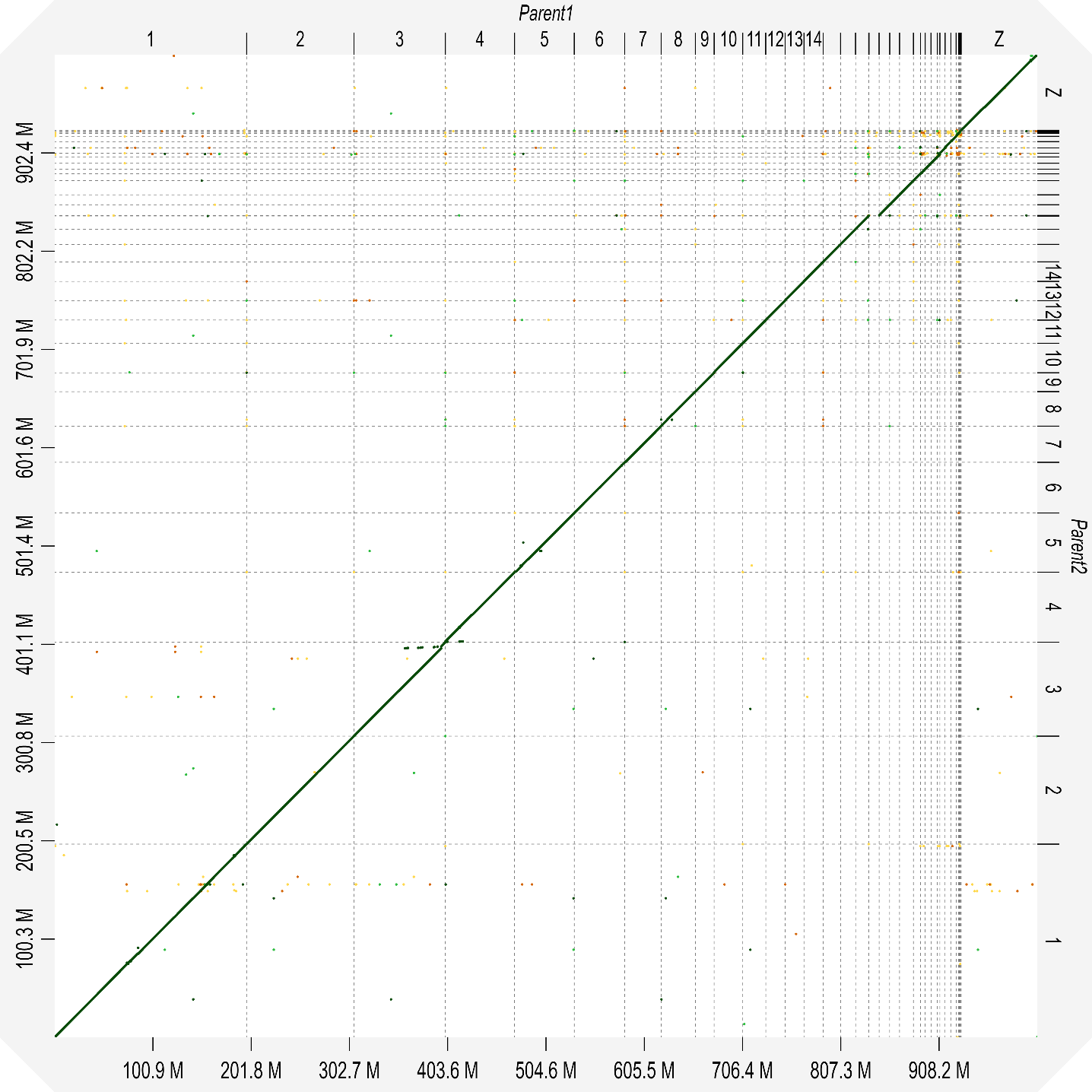

Figure S2 Parent 1 vs. parent 2 alignment

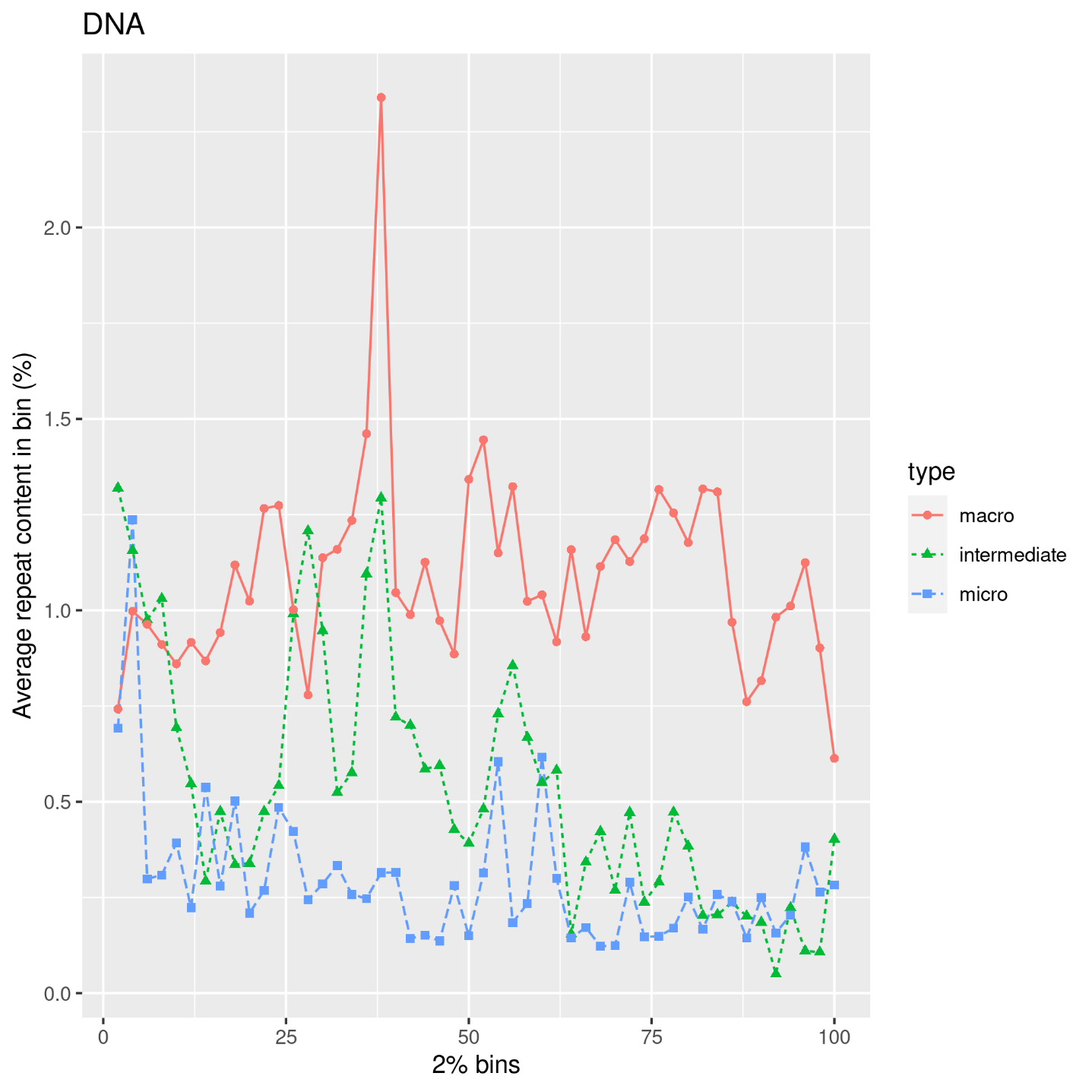

Figure S3: Average DNA repeat content along the chromosomes for macro, intermediate and microcromosomes.

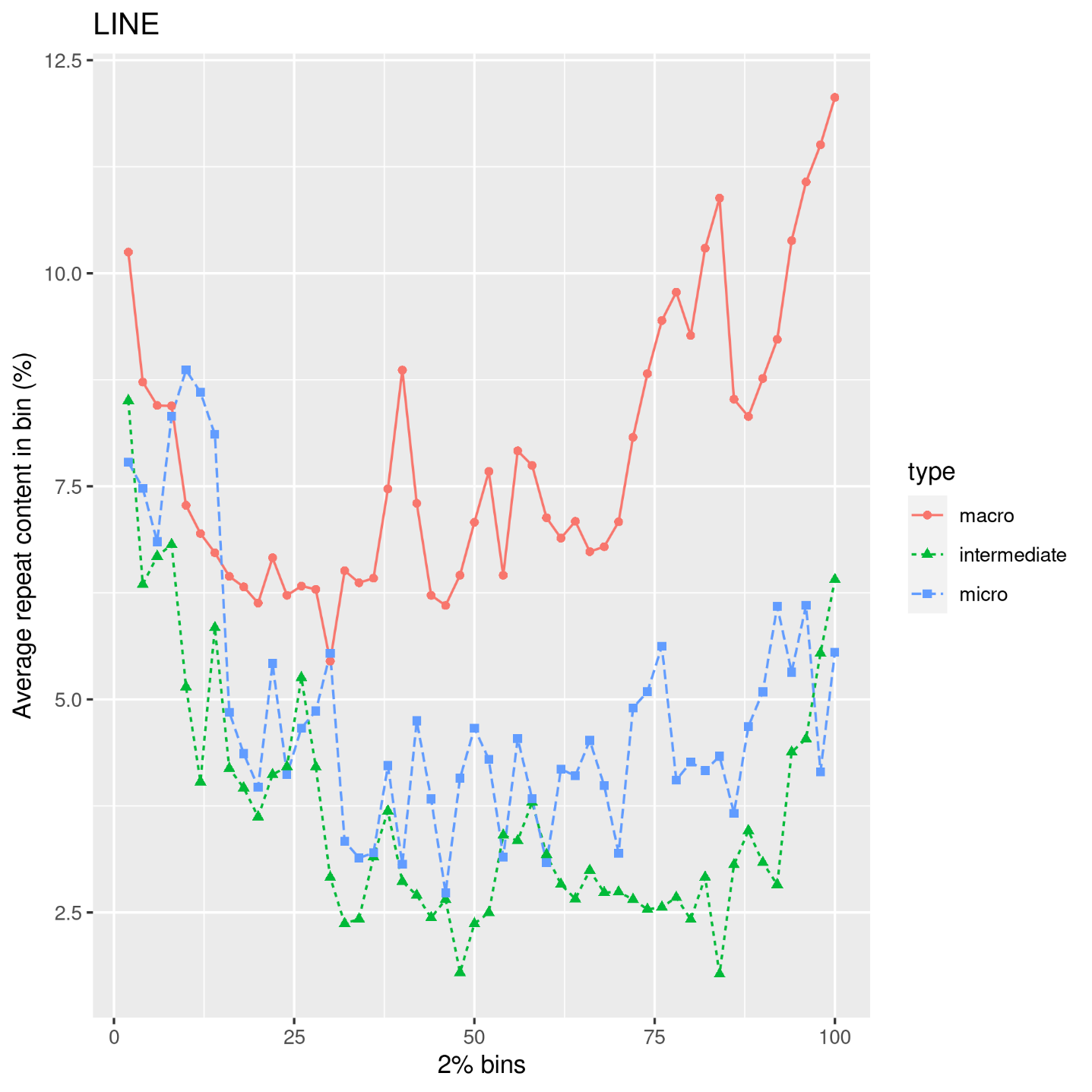

Figure S4: Average LINE repeat content along the chromosomes for macro, intermediate and microcromosomes.

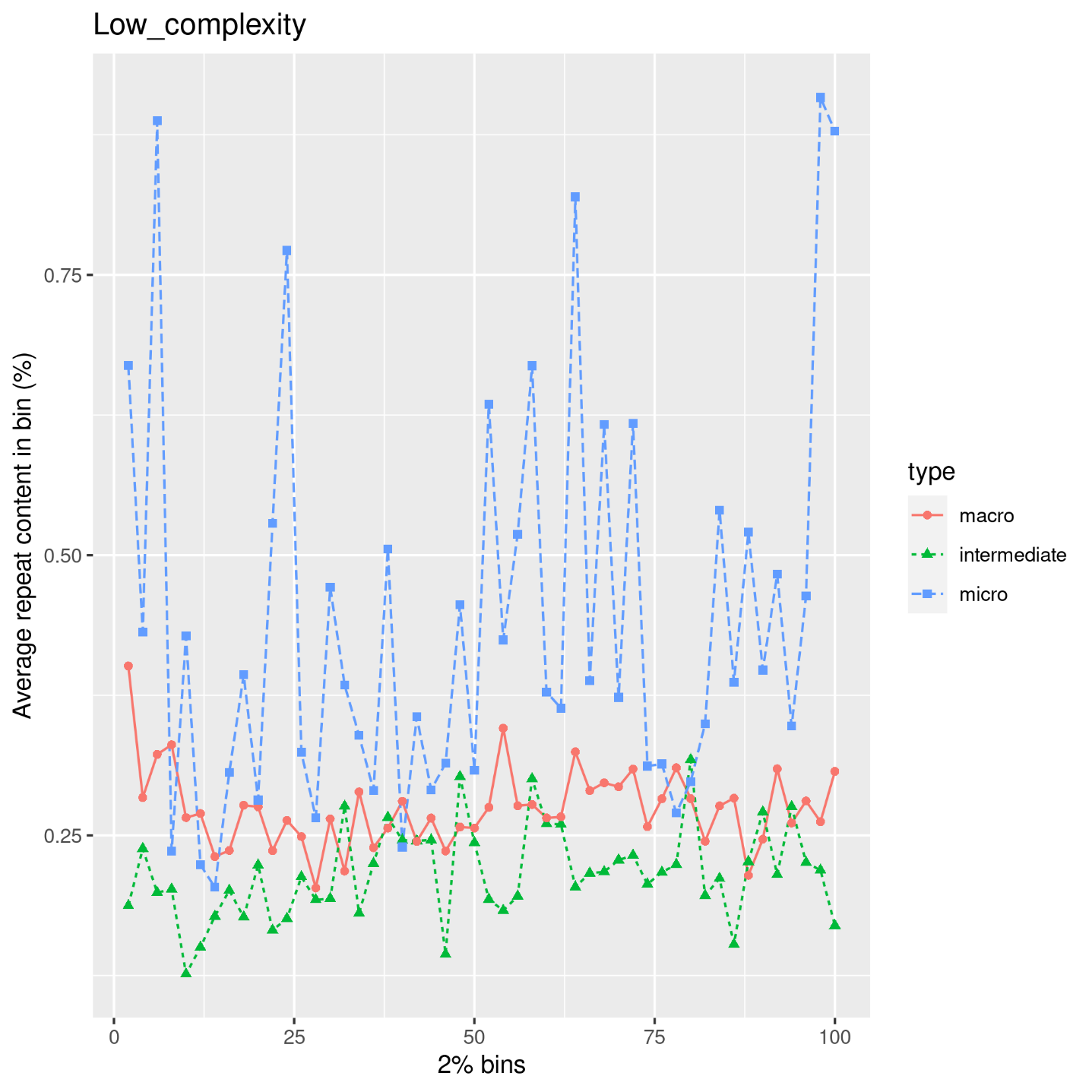

Figure S5: Average low complexity repeat content along the chromosomes for macro, intermediate and microcromosomes.

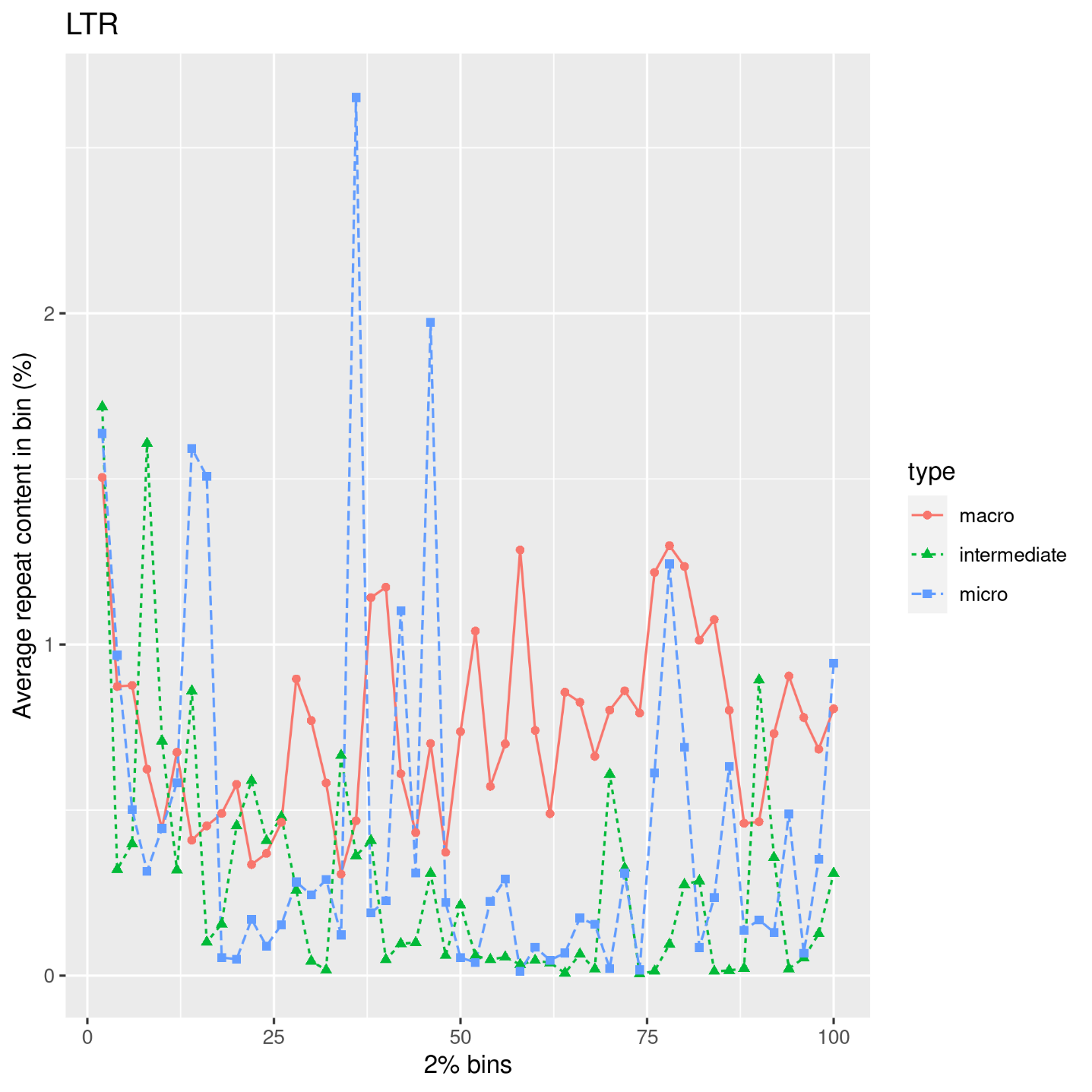

Figure S6: Average LTR repeat content along the chromosomes for macro, intermediate and microcromosomes.

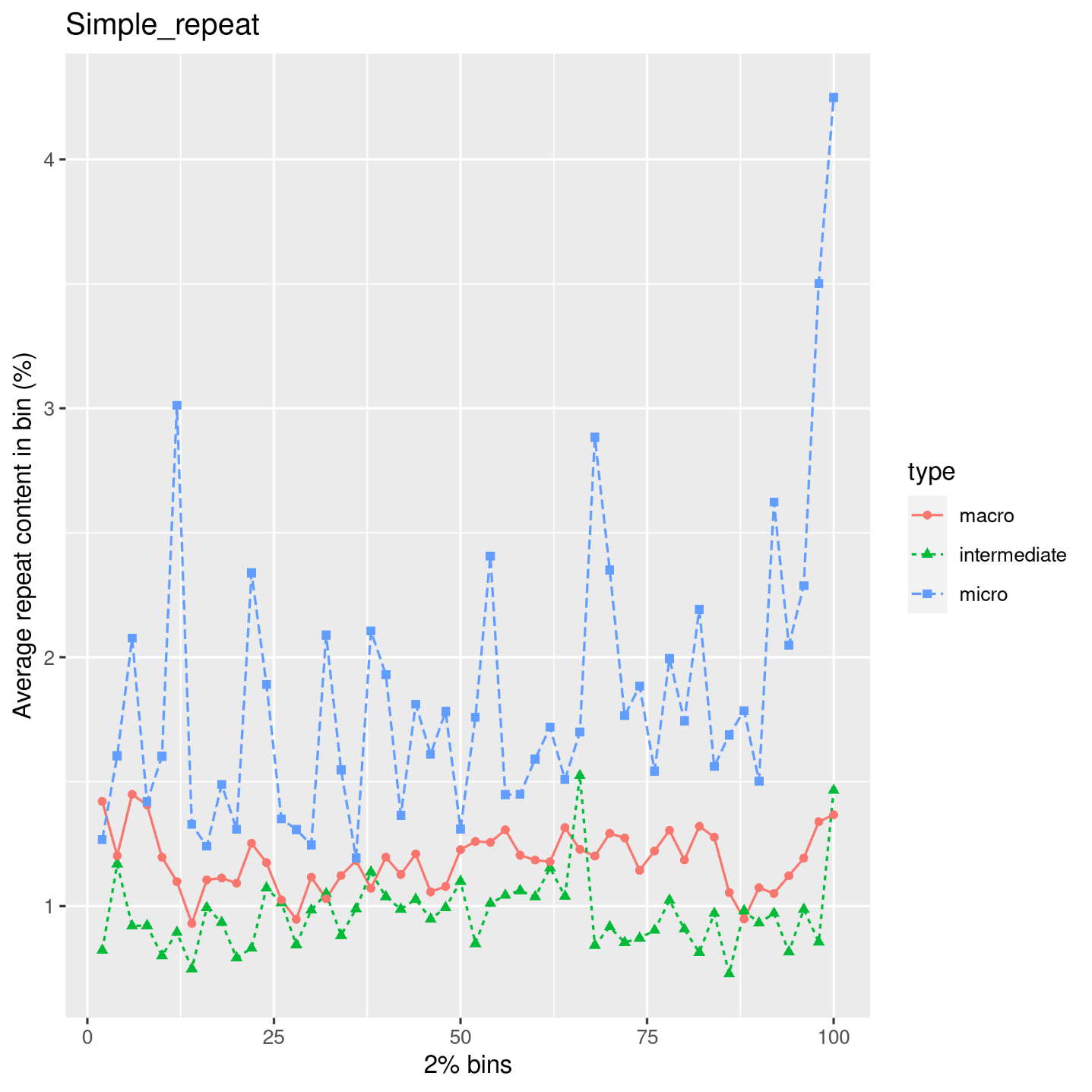

Figure S7: Average simple repeat content along the chromosomes for macro, intermediate and microcromosomes.

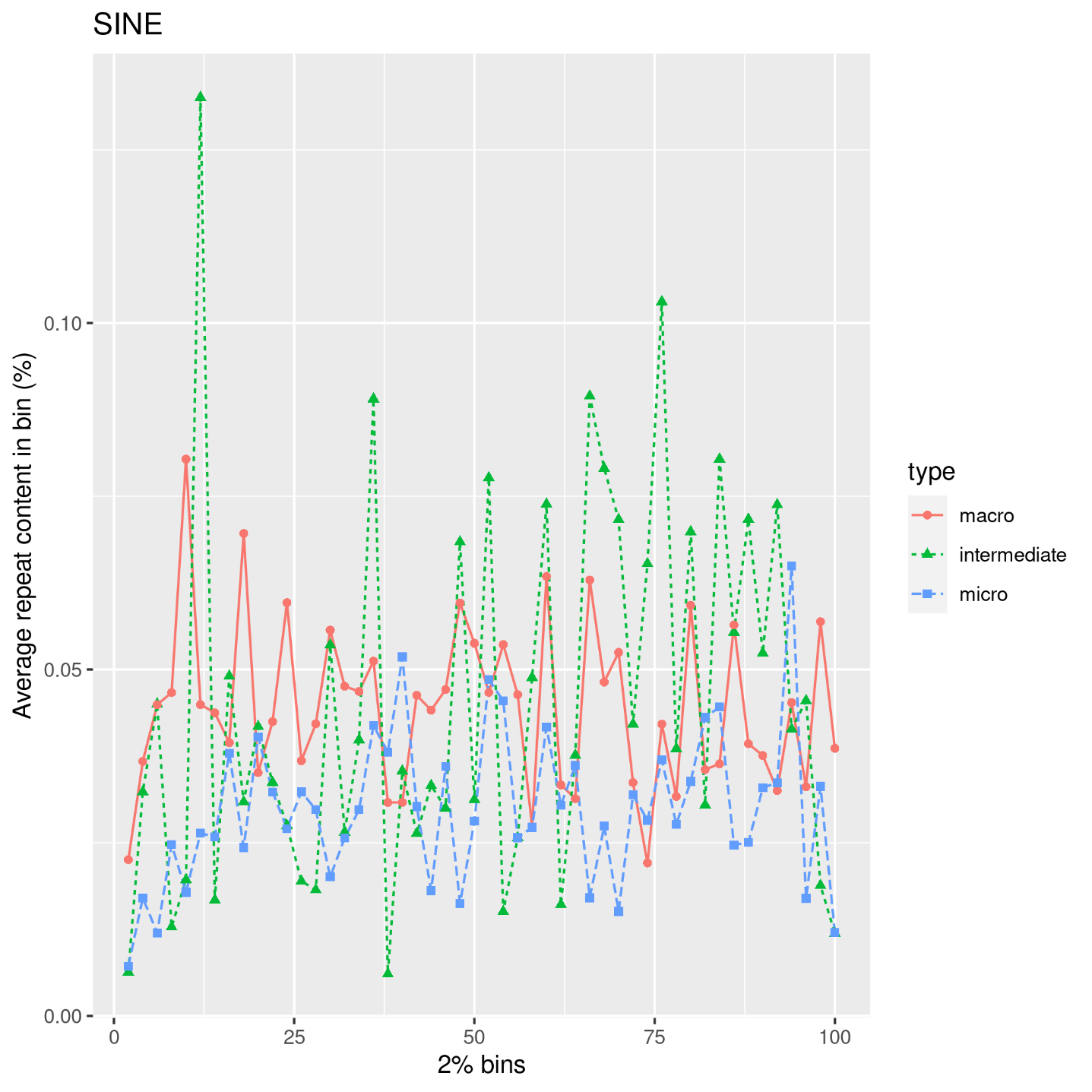

Figure S8: Average SINE repeat content along the chromosomes for macro, intermediate and microcromosomes.

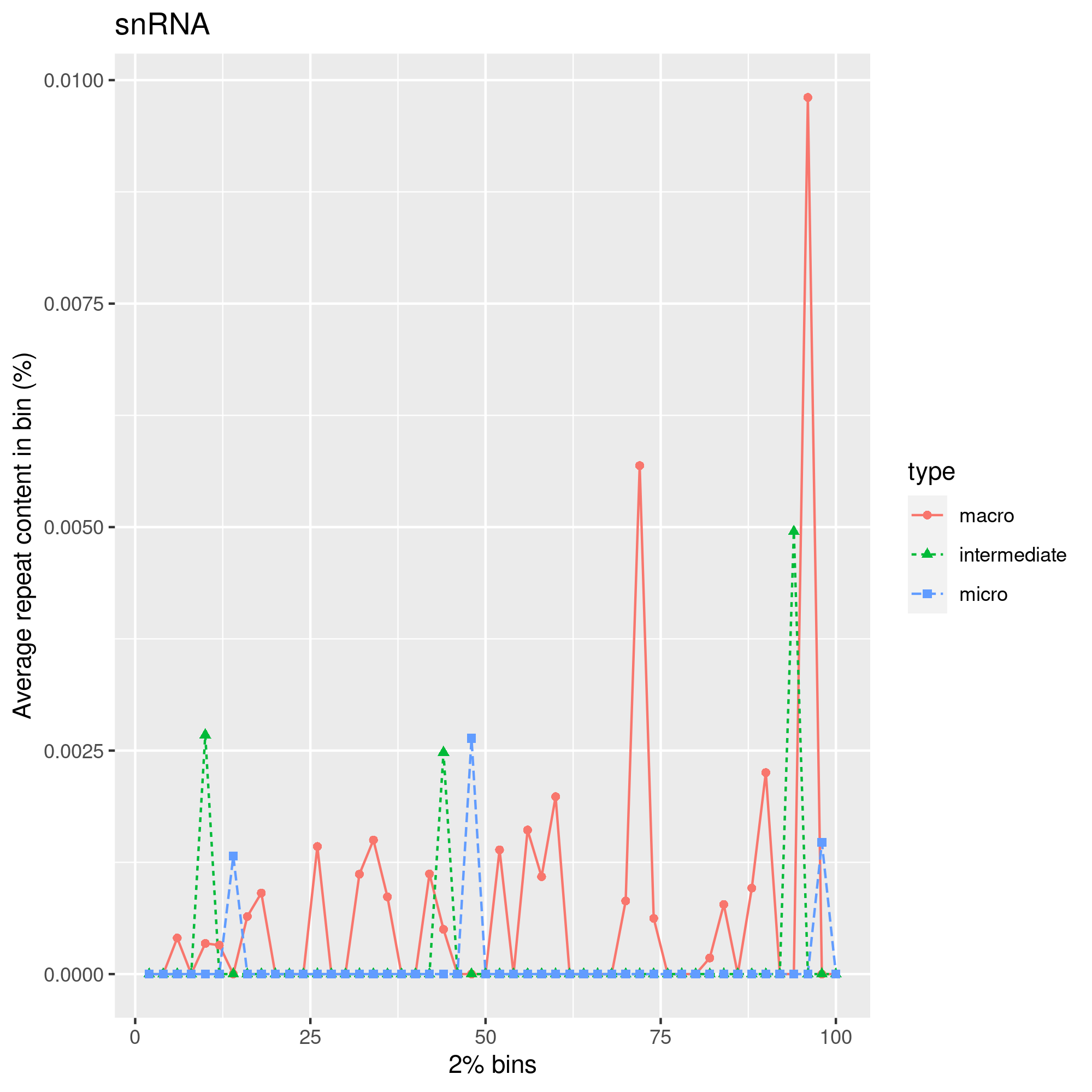

Figure S9: Average snRNA repeat content along the chromosomes for macro, intermediate and microcromosomes.

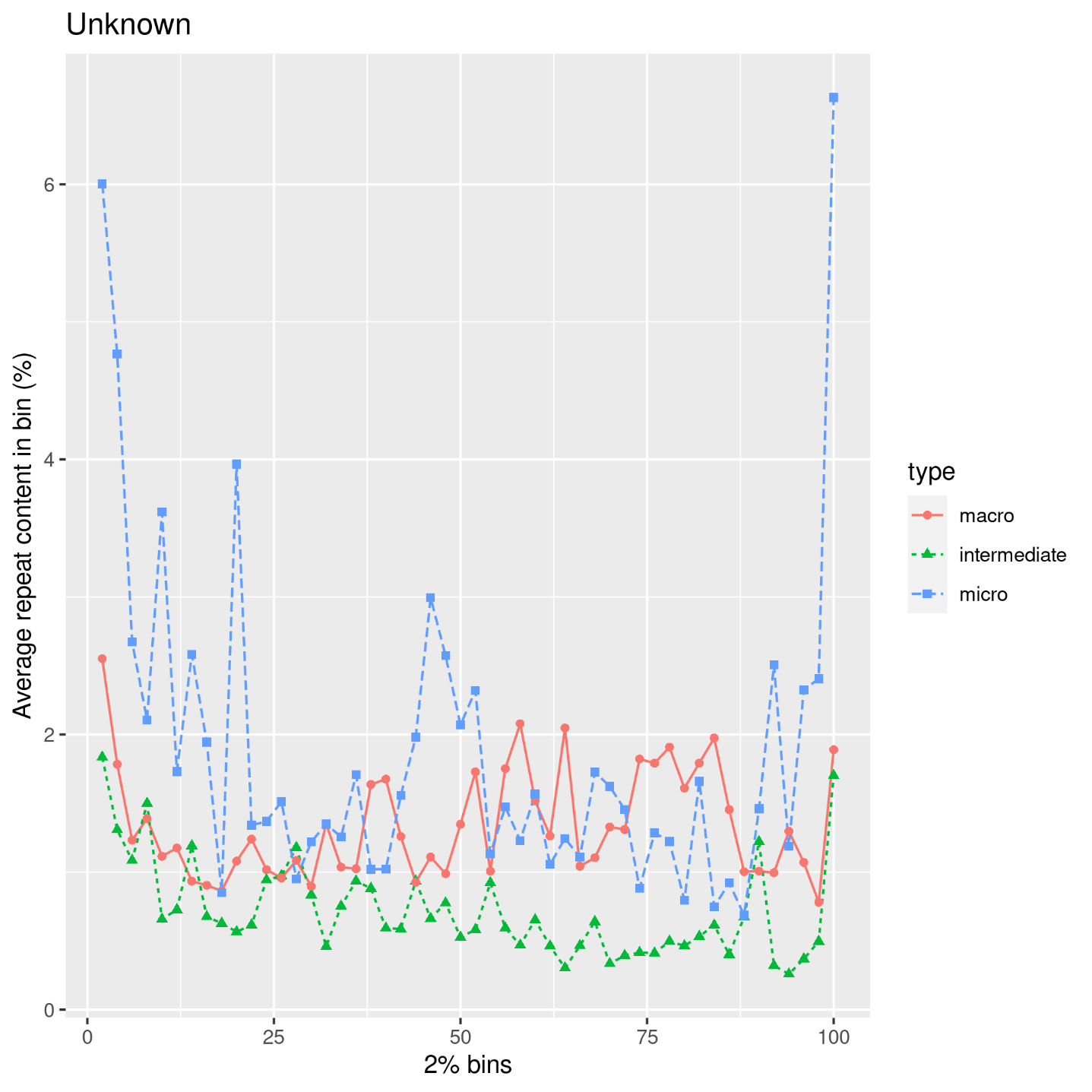

Figure S10: Average unknown repeat content along the chromosomes for macro, intermediate and microcromosomes.

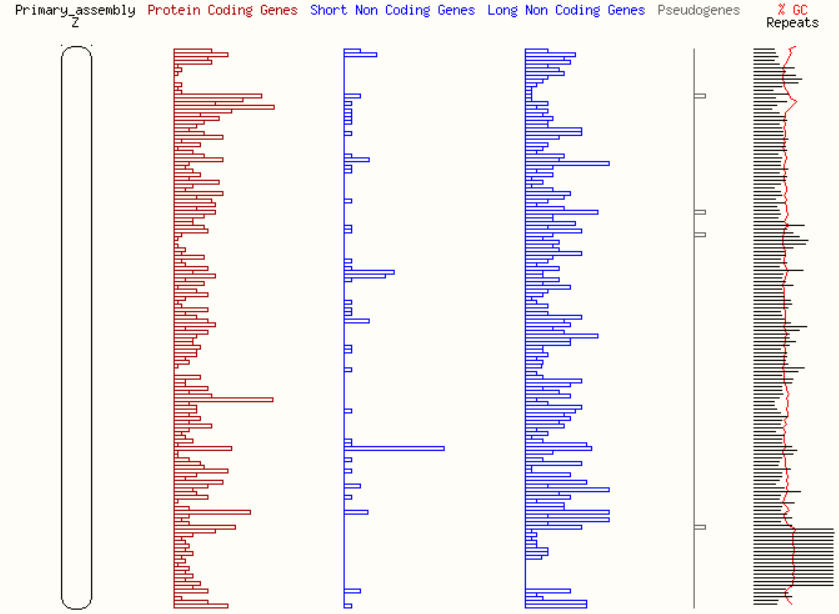

**Figure S11:** **Schematic view of Gal7b chromosome Z and representation of several biotypes of genes and genomic features** (Ensembl, rapid release 15^th^ June 2022, accessed on 27^th^ June 2022).
