## Supplementary figures and images for "A new haplotype-resolved turkey genome to enable turkey genetics and genomics research"

### SupplementaryFile3.pdf

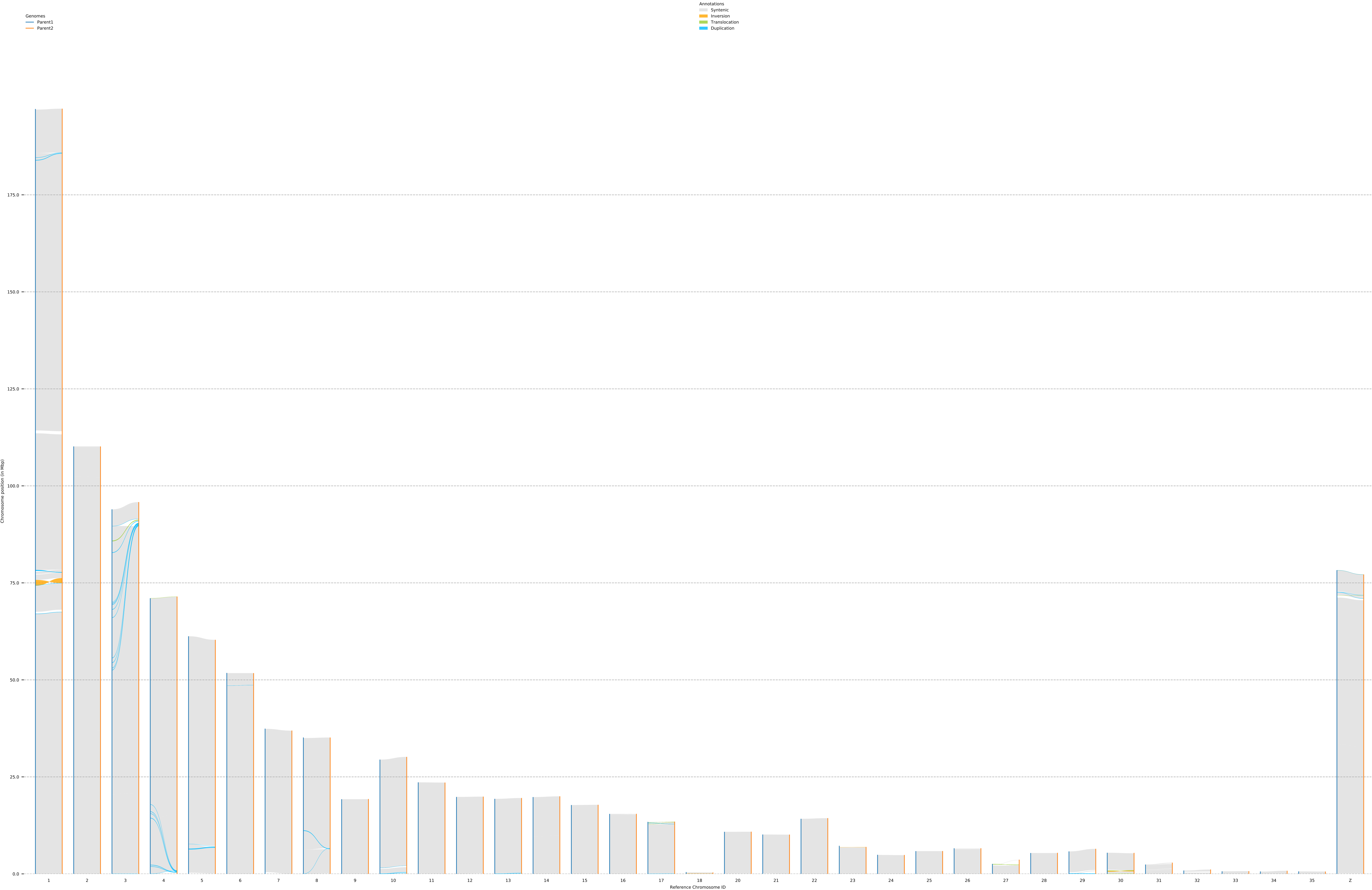
